# Endothelial regulatory circuits control cranial neural crest migration and plasticity

**DOI:** 10.1101/2024.09.17.613303

**Authors:** Zhiyuan Hu, Sarah Mayes, Weixu Wang, José M. Santos-Pereira, Fabian Theis, Tatjana Sauka-Spengler

**Affiliations:** MRC Weatherall Institute of Molecular Medicine, Radcliffe Department of Medicine, University of Oxford; Oxford, United Kingdom; Medical Research Institute, Frontier Science Center for Immunology and Metabolism, Wuhan University; Wuhan, China; Institute of Computational Biology, Helmholtz Center Munich; Munich, Germany; Stowers Institute for Medical Research; Kansas City, MO, USA; MRC WIMM Centre for Computational Biology, MRC Weatherall Institute of Molecular Medicine, University of Oxford, Oxford, United Kingdom

**Keywords:** Cranial neural crest, ETS transcription factors, neural crest migration, neural crest plasticity, epithelial-to-mesenchymal transition, gene regulatory network, spatial transcriptomics, single-cell multi-omics, in silico perturbation

## Abstract

The cranial neural crest (NC) is a migratory embryonic population ideal for studying cell plasticity, motility, and fate establishment. Although NC migration has been linked to changes in cell adhesion, polarity, and signaling, the gene regulatory circuitry governing these processes remained obscure. Using time-resolved single-cell multi-omics, spatial transcriptomics, and gene regulatory network reconstruction, we identified ten programs underlying 23 NC cell states and three spatial trajectories. Using in silico perturbation and systematic CRISPR/Cas9-mediated Perturb-seq, we uncovered novel lineage drivers and an endothelial-like program controlling NC migration, distinct from the epithelial-to-mesenchymal transition (EMT) program. We show that endothelial-like regulons (*fli1a, elk3*) drive migration through direct or tiered activation via the “FOX:ETS-Ebf3a*-*targets” axis, while ETS suppressors (*erf*, *erfl3*) maintain cell plasticity. Using the newly developed SyncReg tool, we identify functional redundancy among ETS regulons, which has thus far obscured their critical roles in NC migration, and we quantify their synergy with retinoic acid receptors, also essential for this process. Our GRN model, combined with novel velocity-embedded simulations, accurately predicted the functions of all major regulons, which were confirmed by in vivo functional perturbations. This study provides a comprehensive, validated cranial NC regulatory landscape, resolving heterogeneous regulatory circuits underlying NC cell motility and plasticity.

## Introduction

The cranial neural crest (NC) is a transient structure in vertebrate embryos that arises from the neural ectoderm but contributes to diverse tissues, including mesenchyme, which originates from the mesoderm in most cases^1^. Cranial NC cells form critical tissues such as cartilage, bone, and connective tissues as well as neurons and glia of peripheral nervous system^2–4^. Defects in cranial NC cells underlie congenital disorders, including Treacher Collins syndrome^5^, craniosynostosis^6^, and other craniofacial abnormalities^7,8^. NC cell multipotency is also implicated in cancers such as melanoma^9^ and Ewing sarcoma^10^. These biological and pathological roles underscore the importance of understanding regulatory mechanisms driving cranial NC development.

Cranial NC migration serves as an ideal model for studying cellular plasticity and motility. Although previous research identified key players^11^, including epithelial-to-mesenchymal transition (EMT) transcription factors (TFs) such as Snail1/2, Zeb2, Twist1^12^, and migratory molecules^13^ such as Small GTPase (RhoA^14^, Rac1^15^), adhesion molecule (e.g., Integrin^16^, Cadherins^17^), and signaling pathways (e.g., PDGF/PDGFR^18^, Cxcl12/Cxcr4^19^), intrinsic epigenetic and transcriptional programs modulating these migratory molecules are poorly understood^20^. Furthermore, it remains controversial which programs drive cell motility in general settings such as tumor metastasis, as complete EMT is rarely found in clinical specimens^20^, leaving the connection between development and tumor cell motility unclear.

Despite significant advances in understanding cranial NC multipotency^21^, fate decision models^22^, and regulatory mechanisms^23^, two key challenges in studying gene regulatory programs remain. First, previous studies have predominantly focused on NC-lineage-specific TFs like AP-2^24^ and SOXE^25^, with limited exploration of TF cooperativity^23,26,27^, including redundancy and synergy. Second, transcriptional regulatory heterogeneity within cranial NC lineages remains poorly understood at single-cell resolution^28–31^. These challenges arise from the nature of cranial NC populations, including their scarce representation at the whole-embryo scale^28–31^, extensive spatiotemporal heterogeneity^32^, and rapid state transitions during development^32^. Current regulatory programs provide a broad-brush overview, averaging regulatory interactions across cell states and fates. This identifies only shared circuits controlled by known *trans-* or *cis*-regulatory elements. However, the full regulatory landscape, a detailed mapping of all subnetworks and systematic perturbation of cell state and fate drivers across developmental stage and axial levels, is still missing. These factors hinder a comprehensive understanding of both normal cranial NC development and disease-related dysfunction.

Here, to fill the gap, we present a comprehensive view of single-cell trajectories and GRN for zebrafish cranial mesenchymal NC development, by leveraging recent advances in single-cell multi-omics, gene regulatory network (GRN) inference^33–35^, precise reverse-genetics perturbations^36–38^, spatial transcriptomics^39,40^, and deep learning algorithms^41,42^. Our GRN analysis identifies ten regulatory programs, revealing lineage-specific time-sensitive roles for 186 regulons. Unlike qualitative models^11^, our GRN framework enables in silico perturbation (RegVelo^43^) to screen TF functions, validated through in vivo Perturb-seq of all major drivers. This approach highlights underexplored TFs, such as those from the EBF, BACH, and TEAD families, quantifies TF functional convergence, and identifies an ETS-mediated endothelial-like program critical for NC migration. Our study detailed profiling and functional validation of the regulatory circuitry controlling cranial NC trajectories in vivo, allowing the interpretation of multi-omics data and aiding the system-scale understanding of physiology and disease^44^. This further enhances our knowledge of NC biology and highlights potential therapeutic targets for related diseases.

## Results

### Single-cell, spatial multiomic landscape of normal and *foxd3*-loss of function NC

To profile cranial NC cells from their inception to differentiation into distinct cell types, we used 10x Multiome to simultaneously capture transcriptomes (single-nucleus RNA sequencing; snRNA-seq) and chromatin accessibility (single-nucleus ATAC-seq; snATAC-seq) from single nuclei at four combined time ranges during first 8-20 hours of embryogenesis (Figure 1A, Figure S1A-C, Supplementary Table 1). Additionally, we analyzed seven distinct time points with the ultra-deep Smart-seq3 single-cell RNA-seq (scRNA-seq)^43,45^. This dual approach captured continuous cell state changes and linked them to specific developmental stages. We defined cell states as the current transitional condition of a cell and cell fates as its developmental path or outcome. Our single-cell multi-omic dataset revealed 24 major cell populations, including NC cells, *foxd3*-expressing neuromesodermal progenitors, and their derivatives (Figure 1B-C, Figure S1B).

**Figure 1:**
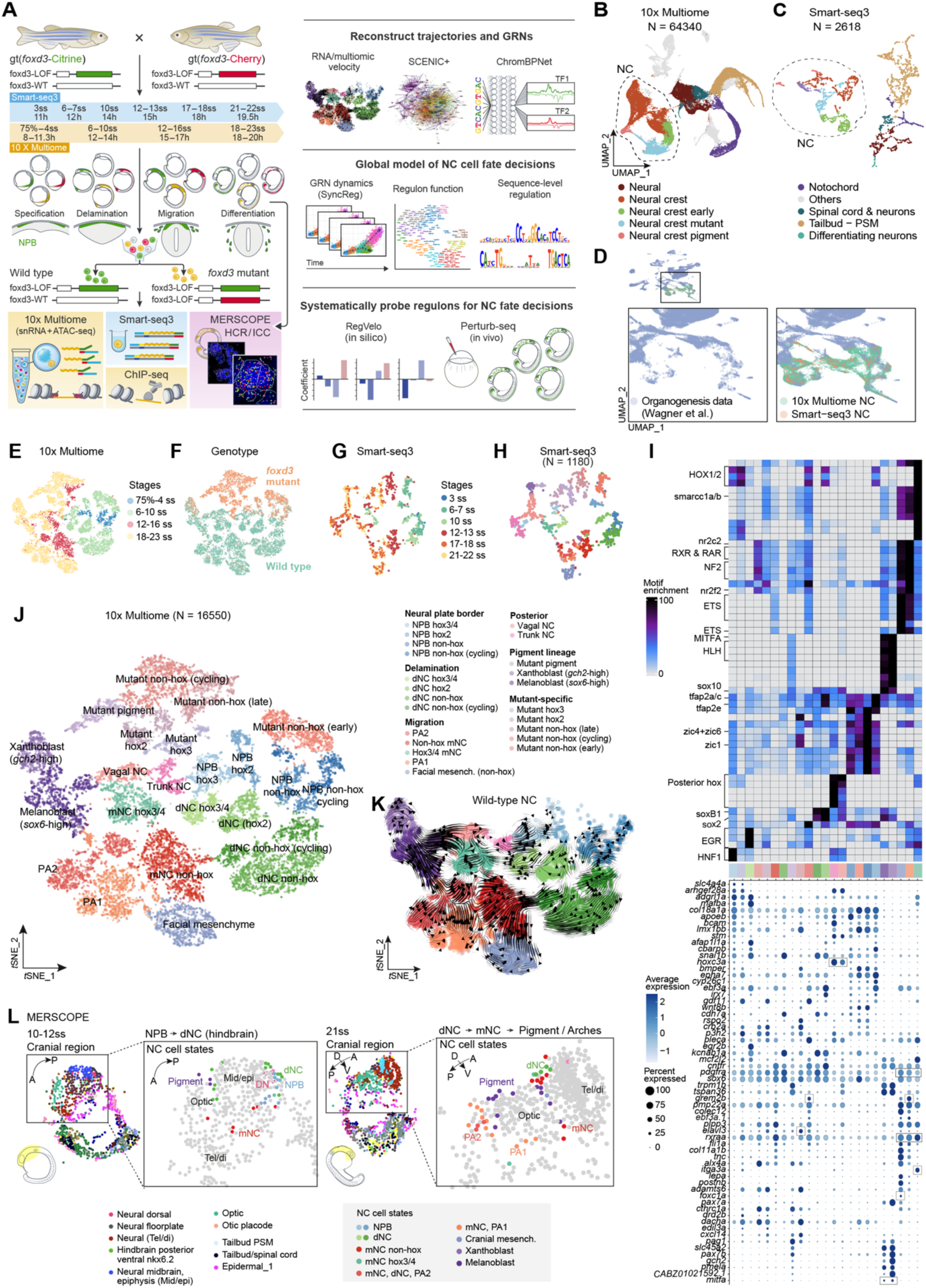
Single-cell multi-omic and spatial transcriptomic profiling reveals the dynamics of zebrafish NC development. **A**, Schematic diagram of the study design. LOF, loss of function; WT, wild type. The Smart-seq3 data were obtained from ref^43^, and other datasets were generated in this study. h, hour post fertilization. **B-C,** Single-cell atlas of zebrafish *foxd3*-expressing NC and posterior populations, shown via uniform manifold approximation and projection (UMAP) visualization of the multiome-snRNA-seq (B) and Smart-seq3 (C) datasets, with individual cells (dots) color-coded by major cell types. PSM, pre-somite mesoderm; NC, neural crest. **D,** Integration of zebrafish organogenesis data^29^ with the multiome-snRNA-seq and Smart-seq3 NC^43^ datasets. Cells (dots) are color-coded by datasets. **E-F**, *t*-distributed Stochastic Neighbor Embedding (*t*-SNE) plots of NC cell atlas in multiome-snRNA-seq datasets, colored by developmental stages (E) and genotypes (F). ss, somites stage; 75%, 75% epiboly. **G-H,** *t*-SNE plots of Smart-seq3 NC dataset, colored by developmental stages (G) and cell states (H). The cell state color panel is the same as (i, j, k). **I,** Motif enrichment (top) and marker gene expression (bottom) in multiome-snATAC-seq and snRNA-seq datasets, against 23 NC cell states (x-axis). **J**, *t*-SNE plot of NC cell atlas in multiome-snRNA-seq datasets, colored by cell states. Mesench., mesenchyme. **K**, RNA velocity analysis results in wild-type NC cells. **L,** MERSCOPE cell state mapping analysis shows the spatial location of cells (dots, colored by broad cell types or NC cell states). A, anterior; P, posterior; D, dorsal; V, ventral; Tel/di, telencephalon/diencephalon; DN, dorsal neural; PSM, pre-somite mesoderm; mesench., mesenchyme; PA, pharyngeal arch.

To explore normal and perturbed NC development, we examined wild-type and mutant embryos lacking the key TF Foxd3^46–48^. Foxd3 is required for craniofacial skeletal differentiation and prevents excessive neuronal differentiation of NCs at the expense of mesenchymal derivatives^49,50^. FoxD3 acts bimodally: as a pioneer factor during NC specification and later as a transcriptional repressor directing NC cells into specific lineages^49^. We isolated wild-type (heterozygous *foxd3*^+/-^) and *foxd3*-mutant (homozygous *foxd3^-/-^*) cells using flow cytometry from embryos of two gene-trap lines, gt(*foxd3*-Citrine) and gt(*foxd3*-mCherry)^50^ (Figure 1A). For the early developmental stage (75% epiboly to 4 somite stage; ss), we combined wild-type and *foxd3*-mutant cells for multiome profiling, and then demultiplexed cellular genotypes using amplicon-seq and machine learning approaches (see Methods; Figure S1D).

We dissected NC ontogeny using single-cell multi-omic and spatial transcriptomic analyses. Unlike prior whole-embryo study^29^, our continuous dataset captured overlooked transitional states of NC development and revealed novel mutant-specific states (Figure 1D)^32^. Clustering NC populations from 8 to 20 hour post fertilization (hpf) across two genotypes yielded 23 distinct cell states, reproducible across 10x Multiome and Smart-seq3 data (Figure 1E-J). Annotated based on developmental stage, these NC states identified in the single-cell data aligned with four key phases of NC development^32^: specification at the NPB, EMT from the neural tube, migration into the periphery, and differentiation. Based on marker gene expression, and motif enrichment, the identified cell states included prospective NC progenitors, delaminating NC (dNC), migratory NC (mNC), and differentiated states at various axial levels such as non-hox facial mesenchyme, pharyngeal arches (PAs), pigment (melanoblasts, xanthoblasts) and progenitors for perspective vagal and trunk NC lineages (Figure 1I-K, Figure S1E, Supplementary Table 2).

To validate NC states and their spatial distribution, we used a customized 300-gene MERSCOPE panel at three developmental stages (Figure S1F-G). Integration with multiome-snRNA-seq clusters confirmed all major NC states, including NPB, dNC, mNC, pigment, non-hox mesenchyme, and PAs and revealed their spatial architecture including NC migratory states along the forebrain-optic and optic-midbrain boundaries and delamination states near the dorsal neural tube (Figure 1L). Our integrated single-cell multiomic and spatial transcriptomic profiling offers a comprehensive, spatiotemporal view for cranial NC.

### Single-cell trajectory analysis depicts contiguous delamination and migration events

To chart NC developmental lineages, we inferred robust trajectories using RNA velocity^51,52^ from multiome-snRNA-seq and Smart-seq3 datasets. This approach predicts lineage dynamics based on the ratios of spliced to unspliced RNA (Figure 1K, 2A-B, Figure S2A). Consistency between latent time and real-world time, coupled with high velocity confidence values, confirmed the reliability of our trajectory analyses (Figure S2B-C). Trajectory analysis of wild-type NC cells linked states from the NPB to NC delamination, migration, and differentiation into various derivative structures. These trajectories precisely reconstructed three migratory routes along the cranial-caudal axis: non-hox (midbrain and 1^st^ pharyngeal arch NC, PA1), hox2 (2^nd^ pharyngeal arch, PA2), and hox3/4 (post-otic NC) (Figure 2A). Whole-embryo HCR validated major cranial NC states, including migratory NC cells (*rxraa*), non-hox facial mesenchyme (*postnb*, *foxc1a* as skeletogenic and cartilage markers^53^), pigment lineages (*mitfa*^54^), pharyngeal arches (*dlx2a*^55^, *grem2b*^56^), and post-otic migratory NC cells (*itga3a*) (Figure 2C, Figure S2D). These validated trajectories recapitulated cranial NC ontogeny at all axial levels.

**Figure 2:**
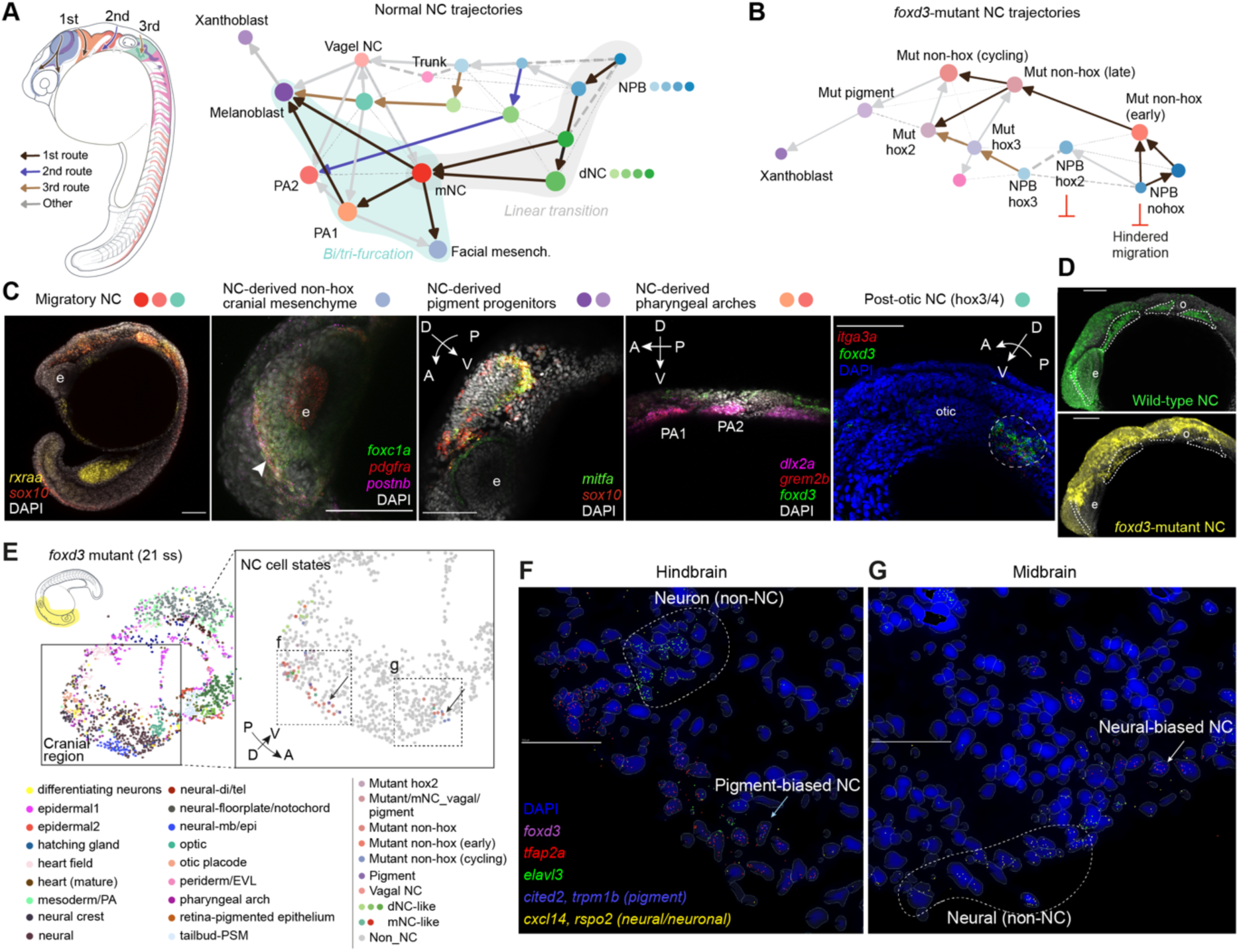
Developmental trajectories and the gene regulatory network (GRN) of zebrafish cranial NC. **A**, Developmental trajectories of wild-type NC reconstructed using multiome-snRNA-seq data with cell states annotated (NPB, neural plate border; facial mesench., facial mesenchyme; mNC, migratory NC). **B,** Developmental trajectories of *foxd3*-mutant NC cells using multiome-snRNA-seq data, highlighting the impact of *foxd3* deficiency on NC development. Mut, mutant. **C,** HCR and IF-HCR images validate the NC states within the whole and cranial regions of wild-type embryos. **D,** Comparative images illustrating hindered NC migration in *foxd3* mutants versus wild type. The green fluorescence (Citrine-positive/mCherry-negative) indicates wild-type NC cells and derivatives, while the yellow (Citrine-positive/mCherry-positive) indicates *foxd3*-mutant NC cells and derivatives. The *foxd3*-mutant NC cells did not populate the target regions, denoted by dashed lines. **E,** Spatial representation of MERSCOPE data of a 21-ss *foxd3*-mutant embryo indicating the major mutant NC states. **F-G,** Two zoomed-in regions in the cranial domain elucidate the pigment-biased (*trpm1b*, mutated *foxd3*, *tfap2a*-coexpressing) and the neural-biased (*cxcl14*, *elavl3*, mutated *foxd3*, *tfap2a*-coexpressing) mutant NC cell states in the hindbrain (F) and midbrain regions (G). The arrow indicates the same cell as in (E). Scale bars, 100 µm.

To investigate the effects of *foxd3* loss, we analyzed NC trajectories in *foxd3*-mutant embryos. The trajectory analysis revealed that *foxd3*-mutant NC cells failed to achieve differentiated states found in wild-type NC cells, with early disruptions emerging during the delamination phase (Figure 2B). Both single-cell trajectories and the whole-embryo immunocytochemistry showed that the second migratory route was absent, and cells in the first (midbrain, rhombomeres 1 and 2) and third (rhombomere 4) routes undertook non-canonical migratory patterns (Figure 2B, 2D).

Spatial transcriptomic analysis confirmed that *foxd3-*mutant NC cells failed to differentiate into standard mesenchymal states, including non-hox midbrain facial mesenchyme, pharyngeal arches, and post-otic derivatives. Instead, they established a new branch containing *elavl3*-expressing cells (Cluster Mutant_nohox_late), which preserved neuronal potential (Figure 2E-G). Another *foxd3*-mutant cluster (Mutant_nohox_cycling) exhibited partial mesenchymal identity, expressing *alx4a*^57^ and *adamts6*^58^ (Figure 1I). A proportion of cranial *foxd3*-mutant NC cells left their dorsal neural plate position but did not reach their facial mesenchyme or pharyngeal arch destinations (Figure 2D). Transcriptomic similarity analysis revealed that *foxd3*-mutant clusters resembled neural floorplate and hindbrain roofplate fates more closely than canonical NC derivatives (Figure S2E). Hence, our analysis reveals spatial sites and new transcriptional identities associated with *foxd3*-mutant cell state changes, including a switch to a potential neuronal ground state^50^, at single-cell resolution.

Overall, these findings demonstrate that *foxd3*-mutant cells can initiate delamination and partial migration but lack the migratory capability and plasticity for cranial NC development. We thus sought to identify downstream Foxd3-regulated programs that are essential for executing cranial NC migration and mesenchymal differentiation.

### GRN and deep learning analyses identify intrinsic regulons for complete migration

As delamination alone did not guarantee that NC cells would reach their destination (Figure 2D), we hypothesized that complete migration requires intrinsic regulators in addition to extracellular environmental signals. Although downstream effectors like adhesion molecules are well studied in NC migration^13^, the key intrinsic TFs driving these effectors—regulating their onset, robustness, and transcriptional switches from the GRN standpoint—remain largely unexplored. Knowledge of the intricacies of regulatory interactions can assist targeted genetic screening and therapeutical strategies for disease^44^.

To reconstruct the full regulatory landscape, we applied SCENIC+^33^ to our multiome data, which identifies regulons comprising TFs, *cis-*regulatory elements, and their target genes by combining concurrent snRNA-seq and snATAC-seq. This analysis identified 186 high-quality enhancer-driven regulons associated with 155 TFs underlying NC development (Figure 3A, Supplementary Table 3). To validate the accuracy of this analysis, we identified twist1b(+) and zeb1a(+), associated with NC delamination^59^, and sox10(+), linked to NC-derived pigment fate determination^54^ (Figure 3A), where ‘(+)’ denotes co-expression of TFs with their target genes. Moreover, core NC TFs, *tfap2a* and *foxd3*, previously shown to be essential for NC formation^24,60^, displayed high out-degree centrality, indicating their top upstream positions in the GRN (Figure 3B, Figure S3A).

**Figure 3:**
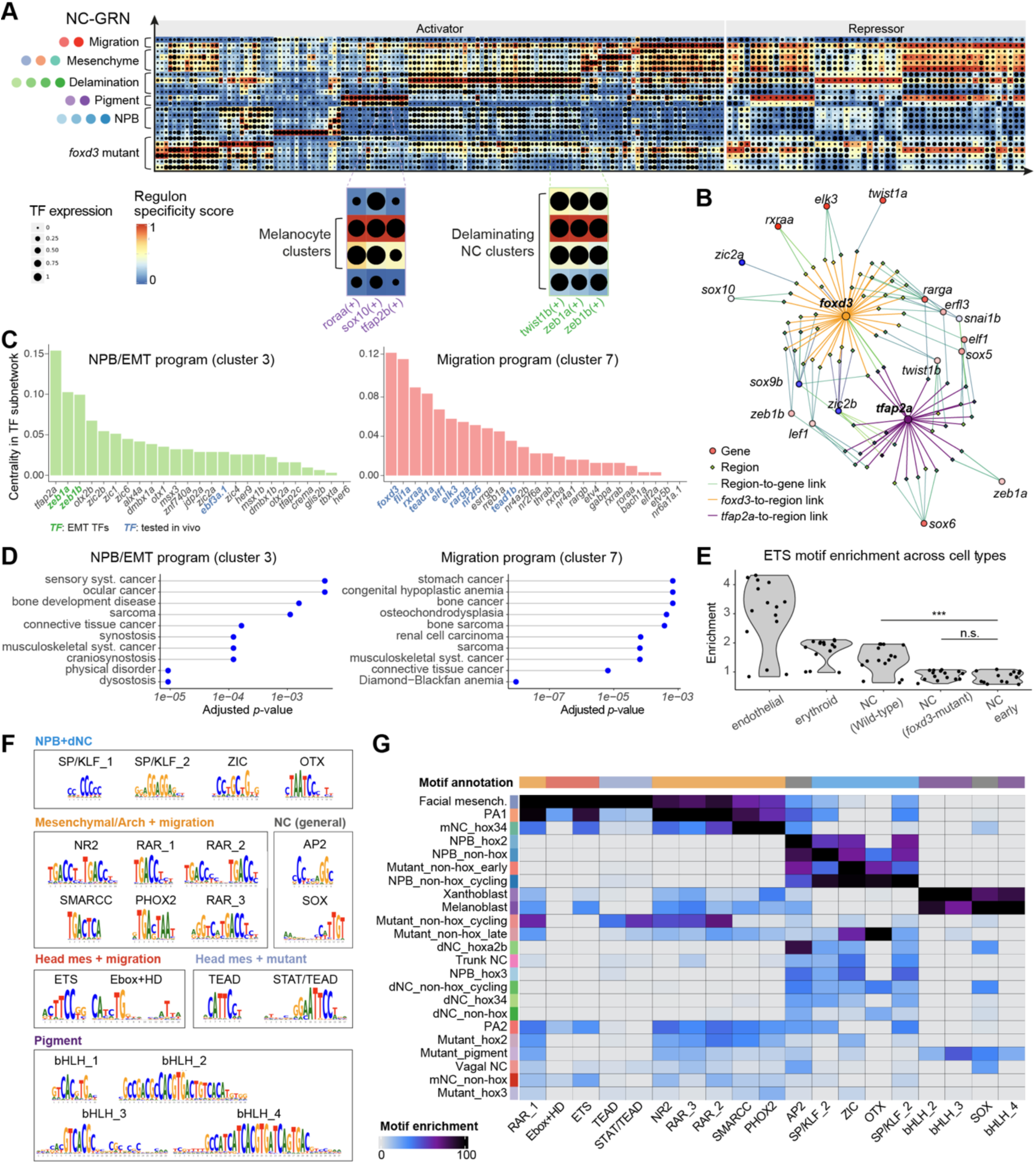
GRN and deep learning analyses identify intrinsic regulons for complete migration. **A**, NC GRN reconstructed from multiome-snRNA-seq and snATAC-seq data, displaying regulon specificity scores (RSS) for regulons (x-axis) across NC cell states (y-axis). The zoom-in panels are examples of six previously recognized regulons. **B,** Network visualizing downstream genes and *cis*-regulatory regions of *foxd3* and *tfap2a*. **C,** TF rankings based on out-degree centrality in the TF-TF subnetwork. Each panel represents a regulatory program, highlighting the TFs (in bold) selected for in vivo Perturb-seq. **D,** Disease enrichment analysis results of putative target genes of these two regulatory programs. **E,** Motif enrichment results of multiome-snATAC-seq data indicate that the ETS motifs are significantly enriched in wild-type NC cells and absent in *foxd3* mutants. **F,** ChromBPNet-derived motifs catalogued by enrichment in cell states. HD, homeodomain. **G,** Motif enrichment analysis across NC cell states. NPB, neural plate border; dNC, delaminating NC; mNC, migratory NC.

We validated SCENIC+-predicted *cis-*regulatory elements using four histone modification profiles: H3K27ac (active enhancers), H3K4me1 (poised enhancers), H3K27me3 (heterochromatin regions), and H3K4me3 (activation of transcription)^61–64^. Based on SCENIC+-inferred GRNs, we identified a group of migratory *cis-*regulatory element candidates that were differentially accessible in migratory wild-type NC cells compared to *foxd3-*mutant NC cells. These migratory elements showed enrichment for H3K27ac (active enhancers) and depletion for H3K27me3 (repression) in the wild-type migratory cells compared to *foxd3-*mutant NC cells (Figure S3B-C). These findings showed that our GRN reconstruction captured the genuine regulatory landscape of migrating NC cells, integrating both transcriptomic and chromatin accessibility data and underscored dynamics of histone modifications, particularly in *cis*-regulatory elements associated with *foxd3*.

Based on regulon activity, we categorized GRNs into ten major programs, including NC specification (e.g., foxd3+, tfap2a+, sox2-), NPB-EMT (e.g., zeb1a+, zic1, zic2b+), migration (e.g., fli1a+, rxraa+), connective tissue and skeletal system differentiation (e.g., ets1+, egr2a+) (Figure 3C, Figure S3D, Supplementary Table 4). To explore the roles of these programs, we performed disease and pathway enrichment analyses with regulon target genes. For the NPB-EMT program, the SCENIC+-inferred target genes were enriched in craniosynostosis (e.g., *AXIN2, ERF, FGFR1/2/3, ZIC1*) and connective tissue cancers (e.g., *BRCA2, SMARCB1, SOX2, SNAI1*) while associating to pathways of negative regulation of neurogenesis (e.g., *notch1a, her5, lfng*) and ephrin receptor signaling (e.g., *efnb1, ephb4a, rasa1a*) (adjusted *p* < 0.0001, by pathway analysis) (Figure 3D, Supplementary Table 4). For the migration program, target genes were enriched in connective tissue cancers (e.g., *FLI1, STAT3, TNC, YAP1, BCL2, CD34*) and pathways associated with cell migration involved in sprouting angiogenesis (e.g., *cxcl12b, fn1a, itga5)* and NC cell migration (e.g., *erbb2, cdh2, foxc1a*) (adjusted *p* < 0.007, by pathway analysis) (Figure 3D, Supplementary Table 4).

The migratory program (Cluster 7) was distinct from the NPB-EMT program (Cluster 3), suggesting that, contrary to previous suggestions where EMT TFs were confounded with pro-migratory roles^65^, intrinsic regulators other than classic EMT TFs mediate NC migration. Top drivers of this migration program, defined by their out-degree centrality in the TF-TF subnetwork, included *foxd3,* ETS TFs (*fli1a, elk3a*), *tead1a,* and retinoic acid signaling receptor TFs (*rxraa*, *rarga*) (Figure 3C). Moreover, ETS motifs (5’-aGGAAg-3’) were enriched in the wild-type migratory and mesenchymal-biased NC cells but diminished in *foxd3* mutants according to the multiome-snATAC-seq data (Figure 3E, Figure S3E), suggesting that *foxd3* may activate ETS TFs by priming them, by directly activating primed enhancers containing ETS motifs, or both. These motifs were also enriched in endothelial and erythroid cells, aligning with early findings^66,67^ and hinting at shared regulatory mechanisms between NC migration and vascular programs.

To further elucidate *cis*-regulatory mechanisms of NC migration using orthogonal computational approaches, we trained the base-resolution deep learning models and conducted de novo motif discovery on our Tn5-bias-corrected ATAC profiles, by applying ChromBPNet^68^ to the multiome-snATAC-seq dataset (Figure 3F). This analysis enabled precise identification of TF binding instances and revealed key motif dominance patterns implicated in NPB, delamination, migration, and differentiation processes. Thus, ETS motifs were associated with mesenchymal fates, RXR/RAR motifs with pharyngeal arch derivatives, and bHLH motifs with pigment lineage differentiation (Figure 3G). These DNA-sequence-based findings corroborated our SCENIC+-inferred regulatory landscapes, reinforcing the roles of ETS, RXR/RAR and bHLH TFs in NC migration and differentiation. Overall, these results showed that an endothelial-like, ETS-enriched regulatory program was activated in migratory NC cells.

### GRN-informed perturbations in vivo and in silico test regulon functions

So far, we have charted the cranial NC molecular lineage and reconstructed the gene regulatory circuits controlling cell fates across a continuous temporal scale. Using this map, we systematically probed regulon functions through global in silico predictions combined with targeted in vivo validations (Figure 4A-B).

**Figure 4:**
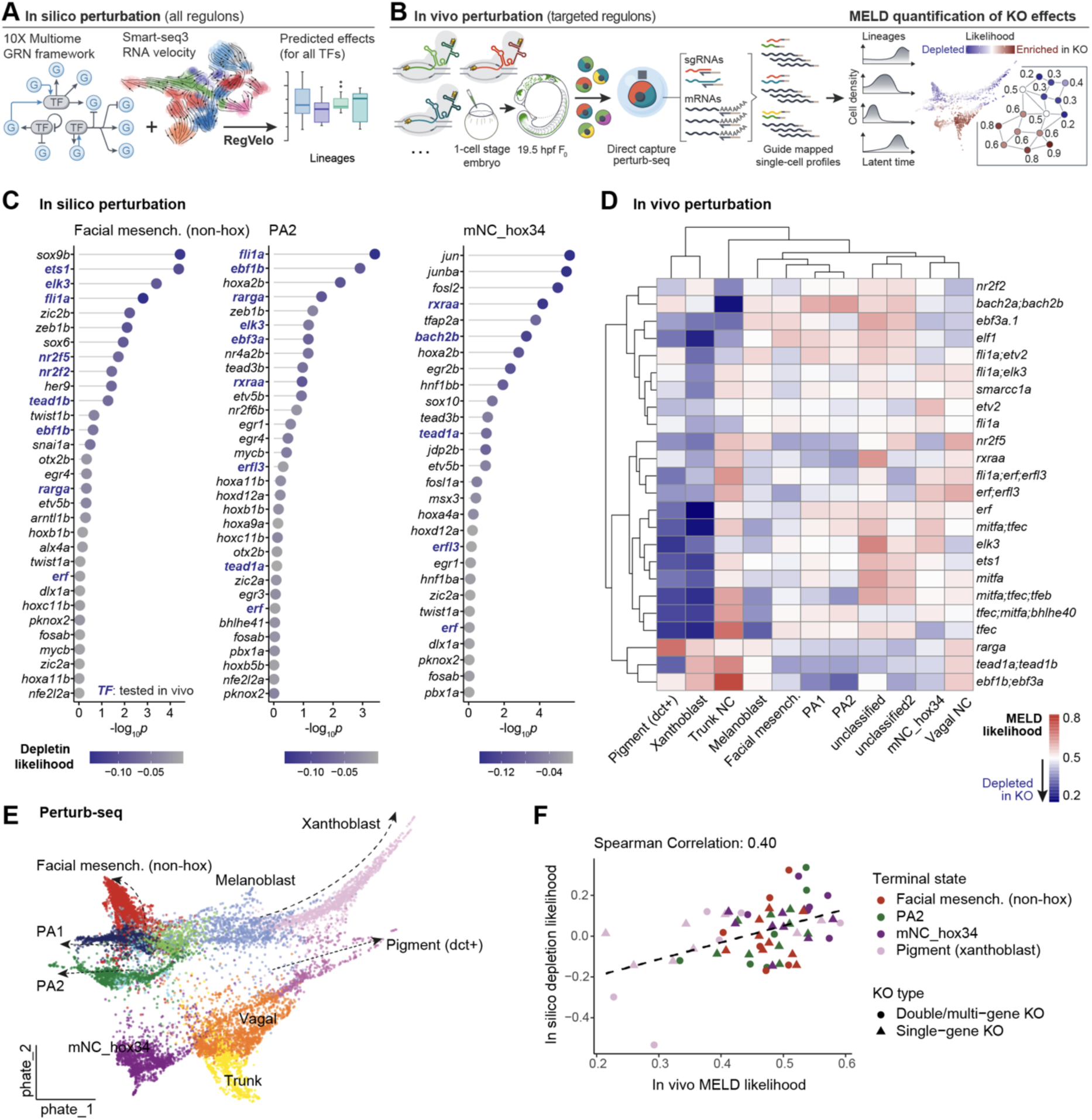
In silico and in vivo perturbation methods probe novel differentiation circuits. **A**, Schematic of RegVelo in silico perturbation methodology. **B,** Diagram outlining in vivo direct-capture Perturb-seq workflow. KO, knockout. **C,** Identification of top TFs through RegVelo in silico simulation for mesenchymal and other cranial lineages, highlighting experimentally tested TFs. All TFs with a negative in-silico depletion likelihood, indicating lineage-specific depletion effects in simulated knockouts, are displayed. **D,** In vivo lineage-specific perturbation likelihood (range: 0-1) across knockout panels and cell states calculated using Perturb-seq data. The in-vivo MELD likelihood lower than 0.5 indicates depletion of the certain lineage, and the likelihood over 0.5 indicates enrichment. **E,** PHATE representation depicting cell states and predicted latent time for all NC cells in the Perturb-seq dataset. **F,** Scatterplot showing the strong correlation between in silico RegVelo prediction and in vivo Perturb-seq likelihood. The dashed line is the linear regression line.

To identify TFs driving cell fate decisions, we conducted the in-silico perturbation using RegVelo^43^ (Figure 4A). RegVelo combines a latent time autoencoder predictor with the non-linear differential equation velocity model, incorporating RNA velocity with a GRN framework to simulate lineage-specific perturbation impacts. Our recent work demonstrated that RegVelo achieved significantly higher accuracy in single-gene and multiple-gene simulation compared to other methods using our NC GRN^43^.

RegVelo simulations identified key drivers of both mesenchymal and pigment fates (*ets1*, *sox6*, *erfl3*, *zic2b*, *ebf1b*), and regulatory diversity among cranial mesenchymal lineages (Figure 4C). For the cranial mesenchymal lineages in midbrain and hindbrain regions, the shared drivers included *rxraa, pknox2*, *tead1a/b* (different ohnologs in different lineages), and *tead3a/b* (different ohnologs in different lineages). For the pre-otic lineages, the shared drivers included *fli1a, elk3, ebf1b, rarga, erf/erfl3* (different ohnologs in different lineages), *zeb1b, egr3/4, otx2b, etv5b, mycb, pbx1a,* and *zic2a.* Individual lineage-specific drivers included *sox6, sox9 and ets1* for midbrain NC with cartilage and mesenchymal fates, *hoxa2b and egr1* for PA2 NC with pharyngeal cartilage and myocardial fate, and *bach2b, egr2b, sox10, her9* for post-otic NC with ceratobranchial, enteric neurons and glial fates. Some TFs (e.g., *ets1*^69^, *sox10*^70^*, sox9b*^25^) are well-known for these roles, but others (e.g., *erf, erfl3, fli1a, elk3*) have not been implicated or characterized in cranial NC development.

To validate these predictions, we performed F_0_ crispant knockouts^36^ in zebrafish, paired with direct-capture Perturb-seq^38^. Our knockout panels targeted 17 newly identified migratory and mesenchymal regulons, focusing on TFs with severe predicted effects using chemically modified single-guide RNAs (sgRNAs) with extended lifespan to ensure robust detection and high knockout efficacy^71^ (Figure S4A-D). To reduce inter-embryo variability, we pooled cells from at least 12 embryos per condition and achieved uniform sequencing depth to minimize biases related to gene coverage (Figure S4E). Because inherent TF redundancy and mosaicism in crispant knockouts may obscure long-term effects, our Perturb-seq analysis workflow was tailored to quantify short-term lineage-specific knockout effects (i.e., a few hours post effective knockout), inspired by CellOracle^34^, using MELD^72^. We dichotomized cells in each lineage based on latent time into early and late lineages, and primarily consider the knockout effects in the late lineages, i.e. the terminal end.

Our Perturb-seq dataset of 17 conditions, combined with another Perturb-seq dataset targeting the pigment drivers^43^, comprised 24 single and combinatorial knockout conditions targeting 23 TFs (Figure 4D-E, Figure S4F-H, Supplementary Table 1). Some targeted TFs (11 out of 22) showed significant downregulation in knockout conditions, supporting effective knockout in these conditions (Figure S4I). In vivo perturbation effects correlated significantly with RegVelo predictions (Spearman correlation coefficients = 0.40), validating the predictive power of our NC GRN (Figure 4F). For example, in vivo Perturb-seq validated that *nr2f5* knockouts significantly depleted non-hox facial mesenchyme lineages, while knockouts of *rxraa*, *rarga*, or *nr2f5* individually affected migratory NC fates in PA2, consistent with RegVelo predictions (*p* < 4×10^-6^, by one-sample one-sided Wilcoxon signed rank exact test, Figure 4D). In contrast, short-term lineage-specific effects from individual ETS TF knockouts (*elk3*, *fli1a*) showed no reduction in non-hox mesenchymal fate, likely due to functional compensation^73^ among ETS TFs (*p* = 1, by one-sample, one-sided Wilcoxon test).

### *erf;erfl3* and *fli1a;elk3* double-TF knockouts disrupt NC migration and differentiation

Next, we performed Perturb-seq analysis in the simultaneous knockout of two or three TFs to investigate the basis of redundancy among ETS TFs.

We prioritized TFs with high regulatory centrality and unexplored roles in lineage determination, selecting *erf;erfl3* and *fli1a;elk3* knockouts because of their predicted disease associations^74,75^, lineage-driving potential, and in silico predicted functions. We selected *erf* and *erfl3* for double knockout as these genes are ohnologs and both are orthologous to human *ERF.* The redundancy among pro-mesenchymal ETS TFs was identified through SCENIC+ GRN analysis, which showed high Jaccard indexes among these TFs (*fli1a*, *elk3*, and *etv2*) based on their shared target genes, as well as the self-regulatory loops and mutual interactions between *fli1a* and *elk3* in the TF-TF subgraph with *fli1a* as the network center (Figure S5A-B).

The *erf;erfl3* knockouts exhibited abnormal craniofacial characteristics potentially associated with craniosynostosis, possibly caused by loss-of-function mutations in *ERF*^75^ (Figure S5C). Perturb-seq analysis demonstrated that *erf;erfl3* knockouts disrupted mesenchymal lineage differentiation, significantly more severely than *erf* single-gene knockouts (Figure 5A-B, Figure S5D). On the other hand, *fli1a;elk3* double knockout depleted the non-hox facial mesenchymal identities more severely than the single-gene knockouts, indicating compensatory roles among pro-mesenchymal ETS TFs (Figure 5C). Similar effects were observed in *fli1a;etv2* double mutants (Figure S5E).

**Figure 5:**
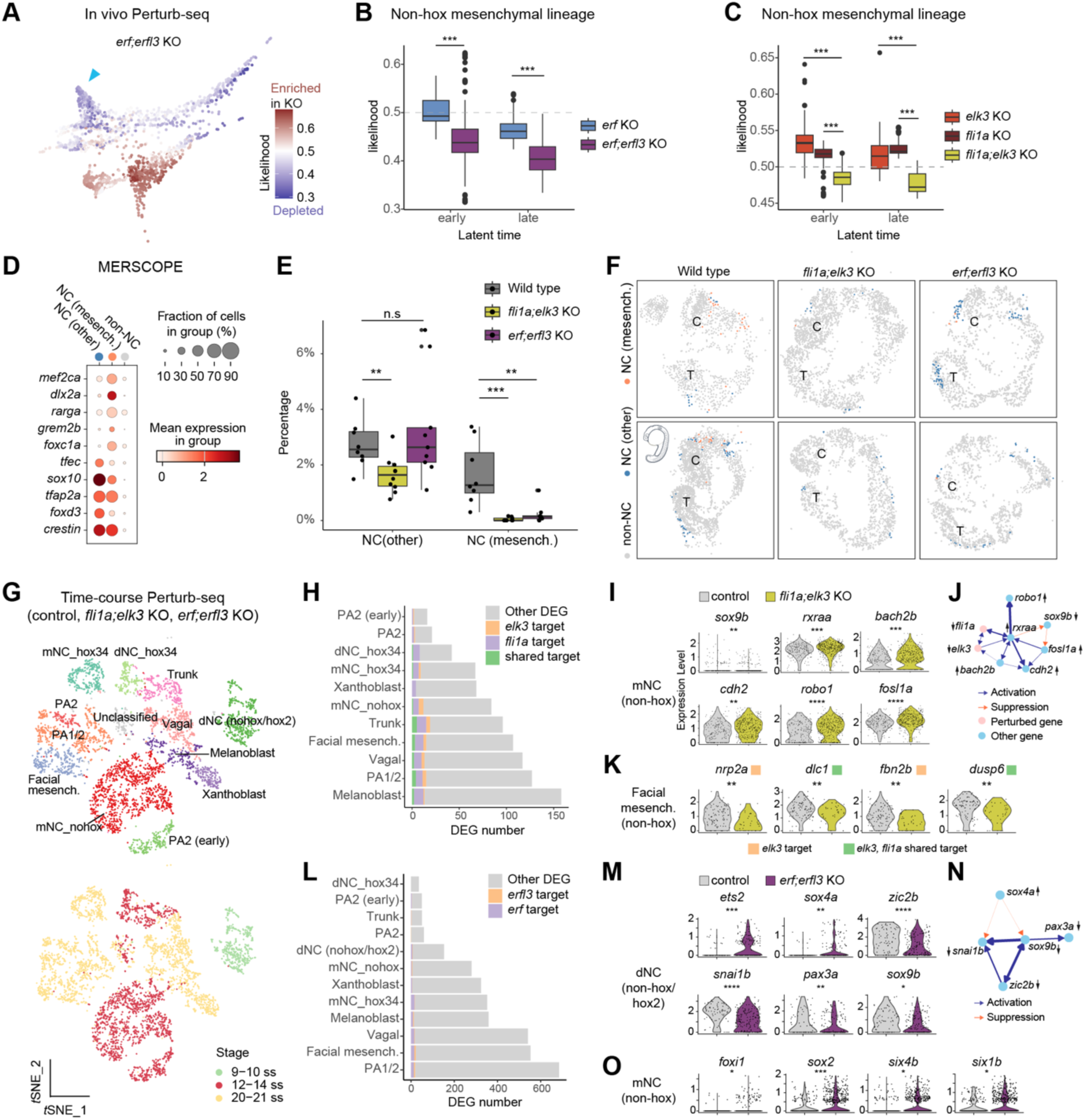
*erf;erfl3* and *fli1a;elk3* double-TF knockouts disrupt NC migration and differentiation. **A,** Perturbation likelihood results highlighting depletion of non-hox head mesenchymal (blue arrow) and pigment lineages in *erf*;*erfl3* double knockouts. **B-C**, In vivo Perturb-seq analysis shows profound depletion effects for *erf;erfl3* knockouts compared to *erf* (B) or for *fli1a;elk3* knockouts compared to *fli1a* or *elk3* (C) in both early and late non-hox facial mesenchymal lineages. *** indicates *p* < 0.001, by Wilcoxon rank sum test. **D,** Dot plot visualizes the marker gene expression in mesenchyme-biased NC (NC, mesench.), other NC, and non-NC cells. **E,** Spatial transcriptomic analysis reveals the significant depletion of mesenchyme-biased NC populations. ***, *p* < 0.001; **, *p* < 0.01; n.s., not significant, by Wilcoxon rank sum test. **F,** Distribution of NC states in spatial coordinates from wild-type, *fli1a;elk3* knockout, *erf;erfl3* knockout embryos at 21 ss. Each dot represents a cell colored by NC states as in (D). C, cranial region; T, tailbud. **G**, *t*-SNE plots visualize the time-course Perturb-seq dataset. **H,** Distribution of TF target genes in differentially expressed genes (DEGs) between *fli1a;elk3* knockouts and control cross NC cell states. **I,** Differentially expressed genes between control and *fli1a;elk3-*knockout cells in the mNC non-hox cluster. ****, *p* < 0.0001; ***, *p* < 0.001; **, *p* < 0.01; *, *p* < 0.05; n.s., not significant, by edgeR. **J,** SCENIC+-inferred GRN subnetwork of TFs and downstream targets from Figure 5I. Directed edge colors represent the SCENIC+-inferred regulatory relationship, and edge weights represent the TF-to-gene importance timed rho value. The arrows next to gene names indicate up-or down-regulation in *fli1a;elk3* knockouts compared to wild type. **K**, Differentially expressed genes between control and *fli1a;elk3-*knockout cells in the facial mesenchyme (non-hox) cluster. **L,** Distribution of TF target genes in differentially expressed genes (DEGs) between *erf;erfl3* knockouts and control cross NC cell states. **M,** Differentially expressed genes between control and *erf;erfl3-*knockout cells in dNC (non-hox/hox2) cluster. **N,** SCENIC+-inferred GRN subnetwork of TFs from Figure 5m. **O,** Differentially expressed genes between control and *erf;erfl3-*knockout cells in mNC (non-hox).

To orthogonally validate these findings, we analyzed double-mutant embryos using the MERSCOPE platform and neighbor enrichment analysis. The spatial transcriptomics confirmed significant depletion of pro-mesenchymal NC and derivatives in both *erf;erfl3* and *fli1a;elk3* knockouts at 21 ss (Figure 5D-F). Moreover, the dorsal-to-ventral migration in knockout embryos was absent, with reduced NC-derived mesenchymal cells near distal sites, such as eye prominences and olfactory placodes (Figure S5F).

To validate the regulatory circuitry orchestrated by these ETS TFs, identify their downstream targets, and assess the temporal effects of knockout on NC migration and mesenchymal differentiation, we performed time-course Perturb-seq across three key developmental stages: early NC migration in the midbrain (9-10 ss; 13.5-14 hpf), active migration in the midbrain NC and delamination at vagal and trunk levels (12-14 ss; 15-16 hpf), and differentiation into cranial NC derivatives (20-21 ss; 19.5 hpf) (Figure 5g).

Differential expression analysis comparing wild-type and *fli1a;elk3*-knockout NC-derived lineages showed that 22% of *fli1a* targets (73/338) and 18% of *elk3* targets (42/233) were significantly differentially expressed (Figure 5H). Since *fli1a* and *elk3* were expressed upon delamination (6-7 ss), we analyzed their knockout effects in migratory NC cells and mesenchymal lineage from the midbrain and PA1 (Figure S5G).

At the active migration stage (12-14 ss), *sox9b, a* key driver of mesenchymal cell fate^25^, was downregulated in the non-Hox midbrain NC, while *rxraa* (associated with neuronal and ganglia development^76^), its putative upstream TFs *(fosl1a*, *bach2b*), and its SCENIC+-inferred target (*cdh2*, *robo1*) were upregulated (Figure 5I-J). Cdh2 (N-Cadherin^77^) plays a role in neuronal migration^78^ and trunk NC migration^79^, and *robo1* is also essential for trunk NC migration^80^. Based on Perturb-seq and SCENIC+-inferred GRN data, we hypothesize that *rxraa* upregulation during midbrain NC migration is driven by a feed-forward loop^81^ to sense the *fli1a*;*elk3* absence, compensating and partially mitigating knockout effects (Figure 5J).

By the differentiation stages (21-22 ss), migration molecules, which are direct targets of Fli1a and Elk3, were significantly downregulated in the facial mesenchymal NC derivatives, including *nrp2a* (*neuropilin 2a*, a semaphorin signaling receptor^82^), *fbn2b* (*fibrillin 2b*, an extracellular matrix component^83^), *dusp6* (*dual specificity phosphatase 6*, involved in vagal NC migration^84^), and *dlc1* (a Rho GTPase-activating protein driving migration^85^) (Figure 5K). These findings strongly support that Fli1a and Elk3 drive NC migration by direct activating migratory molecules and interacting with retinoic acid receptors for transcriptional regulation.

Statistical analysis comparing *erf;erfl3* knockouts to controls in NC-derived lineages showed significant differentiation in 45.6% of *erf* target genes (31/68) and 33.3% of *erfl3* direct targets (10/30) (Figure 5L). Because *erf* and *erfl3* were expressed as early as the NPB phase, we first analyzed *erf;erfl3* knockouts in delaminating NC cells from the midbrain and PA1 at 9-10 ss (Figure S5G). Knockouts resulted in the upregulation of *ets2* (a known *erf* target^86^), *sox4a* (involved in neuronal maturation^87^), *her6* (a pro-neurogenic TF^88^) and the downregulation of *zic2b* (important for NC specification^89^) (Figure 5M). The upregulation of *sox4a*^87^, a TF with over 160 SCENIC+-predicted negatively regulated direct targets, was the presumable intermediate regulator that led to the significant downregulation of NC specification TFs, including *zic2b*^89^, *snai1b*^90^, *pax3a*^91^, *sox9b*^25^ (Figure 5M-N). Thus, *erf* and *erfl3* likely act as “fate-keepers” in NC specification, restricting the repressive activity of pro-neurogenic factors. In the absence of this regulation in *erf;erfl3* knockouts, pro-neurogenic repressors inhibit NC mesenchymal fates while suppressing NC specification factors.

In migratory NC (midbrain and PA1), TFs associated with placodal fates, including *foxi1*^92^*, sox2*^93^*, six4b*^94^, and *six1b*^95^ were upregulated (Figure 5O). At 20-21 ss, these changes were milder in the facial mesenchymal cluster due to lineage depletion and, thus, we focused on the less-depleted PA1/2 clusters; in knockouts, the downregulated genes included SCENIC+-inferred targets (*kcnn4*, *tgfbr3*^96^, *gse1*^97^), as well as molecules important for cell adhesion, migration, and signaling (e.g., *palld*^98^, *cdh7a*^99^, *itga8*^100^, *pdgfra*^18^, *plxna1a*^101^, *epha2a*^102^, *nrp2b*^82^, *fgfr2*^103^) (adjusted *p* < 0.01, by edgeR; Figure S5H). These results suggest that *erf* and *erfl3* regulate genes involved in extracellular matrix organization, cell adhesion, PDGF signaling^18^, and semaphorin signaling^82^ to drive the migratory and differentiation programs critical for NC cell fate decisions.

Overall, these findings demonstrate that *fli1a;elk3* and *erf;erfl3* are essential for cranial NC migration and pro-mesenchymal differentiation. However, they regulate distinct mechanisms despite sharing similar binding motifs. *Erf* and *erfl3* maintain NC plasticity by preventing pro-neurogenic activation of *sox4a* and *her6*, while *fli1a* and *elk3* drive an endothelial-like program for NC migration by regulating a suite of downstream genes known to effectively mediate migration.

### The endothelial-like program differs from the EMT program

Based on our analysis, we hypothesized that cranial NC migration is driven by an endothelial-like program, separate from the well-recognized EMT regulons such as ZEB and TWIST. To test this hypothesis, we needed a detailed temporal view of regulon activity changes.

We constructed the dynamic model integrating multiple modalities using MultiVelo^35^, which combined transcriptomic and chromatin accessibility data (Figure 6A). Focusing on non-hox trajectories reduced confounding kinetic differences and improved latent time prediction. This multiomic approach outperformed RNA velocity in clearly delineating delamination and migration phases, achieving better alignment with real-world time (Figure S6A-C). Using this model, we predicted cell state transitions along the non-hox trajectory, spanning the NPB, delamination, migration, and differentiation states, using CellRank^104,105^. The terminal state probabilities, representing the likelihood for each differentiated cell state, were low during NPB and delamination phases but increased substantially during migration, indicating the onset of fate bias (Figure 6B).

**Figure 6:**
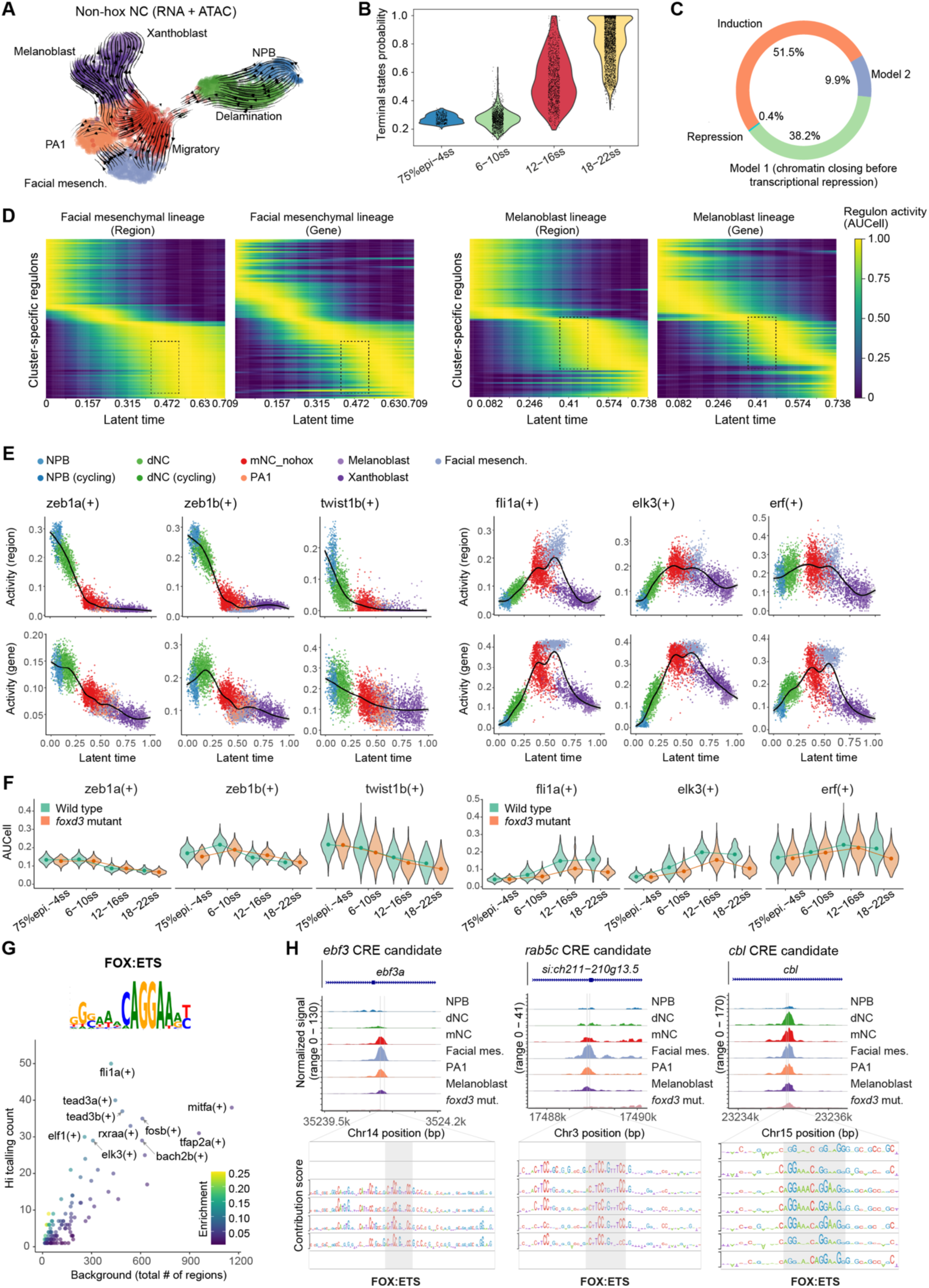
Unraveling EMT and differentiation dynamics through temporal GRN and deep learning analyses. **A**, Multiomic velocity streams of non-hox NC trajectories. **B,** Violin plot demonstrates the relationship between chronological stages and CellRank terminal states probability, which is positively associated with later developmental stages. **C,** Pie plot showing the proportions of genes following specific transcriptional regulatory models (indicated by colors). Models 1 and 2 represent that chromatin closing happens before or after transcriptional repression, respectively. **D,** Heatmaps showing region- and gene-based regulon activity cascades (n = 95) along latent time for facial mesenchymal and melanoblast lineages, with priming events marked by boxes. **E,** Trend plots for representative EMT regulons (*zeb1a*, *zeb1b*, *twist1b*) and ETS regulons (*fli1a, elk3, erf*). The y-axes indicate regulon activity, and the x-axes indicate latent time. **F,** Stage-specific and genotype-specific regulon activities of representative EMT and ETS regulons. The lines link the median of each group. **G,** The de novo discovered FOX:ETS motif (top panel) and its distribution in positively regulated regulons (bottom panel). **H,** Putative target genes *ebf3*, *rab5c*, and *cbl* downstream of the FOX:ETS motif, based on chromatin accessibility of *cis*-regulatory element candidates (track plots, top panels) and the highlighted FOX:ETS TF binding instances (contribution score profiles, bottom panels).

At the gene level, we observed that velocity dynamics predominantly adhered to two models: the induction model (52%), characterized by partial kinetics with only the induction phase, and Model 1 (38%), characterized by full kinetics with both induction and repression phases and chromatin closing before transcriptional repression. This suggests that transcriptional repression was primarily driven by chromatin remodeling in this NC developmental process (Figure 6C). To dissect lineage-specific dynamics leading to differentiation, we incorporated cell fate probabilities into these temporal processes and defined cluster-specific regulons using regulon specificity scores (RSSs). Analysis of regulon activity cascades over latent time uncovered that regulon-wise chromatin accessibility preceded gene expression for pigment and mesenchymal lineage markers, emphasizing enhancer-driven gene activation as a key regulatory mechanism defining NC-derived lineages (Figure 6D).

Using our model of regulatory dynamics, we identified a temporal separation between EMT regulons and the novel endothelial-like NC migration program. EMT is critical for NC delamination and migration^12^. Our analysis of GRN dynamics revealed that EMT-related regulons, including zeb1a(+), zeb1b(+), and twist1b(+), were active as early as the NPB phase in terms of region-based activities, and in terms of target-expression-based activities they became active during delamination (Figure 6E). In contrast, ETS regulons, i.e., fli1a(+) and elk3(+), were primed after EMT initiation and prior to differentiation, suggesting a temporal role different from EMT regulons (Figure 6E). Moreover, erf(+) was activated at region levels during the NPB phase, supporting its fate keeper role distinct from fli1a(+) and elk3(+).

Overall, we demonstrated that our dynamic model built using MultiVelo accurately matched developmental stages and reflected the temporal progression of cell fate decisions. This analysis further confirmed that the ETS-driven endothelial-like program, distinct from the EMT program, controls NC migration.

### *foxd3* activates endothelial-like regulons through FOX:ETS synergy

Given the temporal asynchrony between the EMT and endothelial-like migration programs, we investigated whether both programs were regulated by *foxd3*. Both EMT and ETS regulon activities were significantly reduced in *foxd3* mutants; however, ETS regulons were more severely affected, with fold changes of 4.7%-12.8% for EMT regulons compared to 22.6-30.1% for *fli1a* and *elk3* regulons. We demonstrated that *foxd3* was essential for priming ETS motif-harboring regions in specified NC cells (Figure 3E). However, the precise regulatory mechanism by which *foxd3* drives endothelial-like regulons remained unclear.

Using TFModisco-lite^106^, we identified the FOX:ETS composite motif^107^, which explains the role of *foxd3* in ETS motif priming (Figure 6G). This motif was reported to be involved in development and lymphoma^27^, and is commonly found upstream of endothelial genes^107^. Predicted TF binding sites for this motif were enriched in fli1a(+), elk3(+), tead3a/b(+), fosb(+), tfap2a(+), mitfa(+), elf1(+), and bach2b(+) regulons (Figure 6G). To examine the functions of this motif in NC development, we defined high-confidence target genes by intersecting SCENIC+-inferred region-to-gene links for *fli1a, elk3, ets1, erf, erlf3* regulons with ChromBPNet-predicted TF binding regions. The pathway enrichment analysis showed that these high-confidence downstream targets in NC were enriched in pathways of sprouting angiogenesis (e.g., *sh3pxd2aa, msna, cdh11, ednraa, nfatc2a*) and regulation of MAPK cascade (e.g., *ccny, itgb1b, ldb2a, add3a*) (adjusted *p* < 0.001; Figure S6D), implying that cranial NC exploits the endothelial-like regulatory program for migration.

In addition to priming ETS TF binding sites, our analysis also demonstrated that foxd3 directly regulates the endothelial-like program by transcriptionally activating *elk3* (Figure 3B). The *elk3* expression levels were downregulated in *foxd3* mutants starting from the delamination stage, supporting the upstream regulatory role of Foxd3 (Figure S6E).

The identified FOX:ETS targets included *ebf3a*, *rab5c* (a small GTPase involved in endothelial cell adhesion and migration^108^), and *cbl* (an E3 ubiquitin ligase involved in melanoma cell invasion^108^) (Figure 6H). Among these, *ebf3a* (orthologous to human *EBF1*) was reported to modulate osteoblast and adipocyte lineages^109^. Our Perturb-seq analysis demonstrated that double-TF knockout targeting *ebf3a* and its ohnologue *ebf1b* (*ebf3a*;*ebf1b* KO) resulted in the depletion of all cranial mesenchymal lineages (Figure 4D). Furthermore, *ebf3a* regulon targets were significantly enriched in pathways related to ephrin receptor signaling pathway (e.g., *rasa1a, efna3b*) and GTPase activation (Figure S6F, Supplementary Table 5). Together, these results suggest that the FOX:ETS composite motif orchestrates the secondary steps of NC migration and mesenchymal lineage differentiation by activating ebf3a(+) and other migration-related genes.

Overall, these results refine the dual regulatory roles of Foxd3 in NC development: Foxd3 partially assists EMT-mediated delamination during the early phase and, more importantly, activates the ETS-driven endothelial-like program and related TFs like *rxraa* during the second phase of migration. While *ets1*, an ETS TF, has been implicated in NC migration^69,110^, our findings revealed potential synergistic roles among other ETS TFs in coordinating NC migration and lineage differentiation, which were previously unrecognized. Our work suggests that ETS TFs, well studied in vascular development and tumor migration^69,107,111,112^, also play vital roles in NC migration and fate determination, likely operating through a shared regulatory circuit.

### Novel computational methods identify TFs with convergent functions

TF synergistic control contributes to binding and regulatory specificity^113–116^, which is critical for understanding mutation effects and refining medical interventions^117^. Despite some synergistic interactions that have been previously studied^118,119^, a global understanding of these effects in NC development remains elusive. To address this, we investigated the potential synergistic effects between the endothelial-like programs and other TFs, including members of the RXR/RAR family identified within the same migration program (Figure 3C).

Inspired by the FOX:ETS synergy, we systematically examined TFs with convergent functions contributing to NC migration and differentiation into mesenchymal derivatives. We characterized TF functional convergence using two metrics: redundant effects, defined by target sharing and high motif similarity (e.g., *fli1a* and *elk3*), and non-redundant cooperative effects, involving target sharing but low motif similarity. To take temporal dynamics into account for TF cooperativity, we developed the “regulatory synchronization (SyncReg)” algorithm, which maps regulon temporal activity patterns onto a two-dimensional plane, extracts features by neural networks and computes similarity among regulons (Figure 7A).

**Figure 7:**
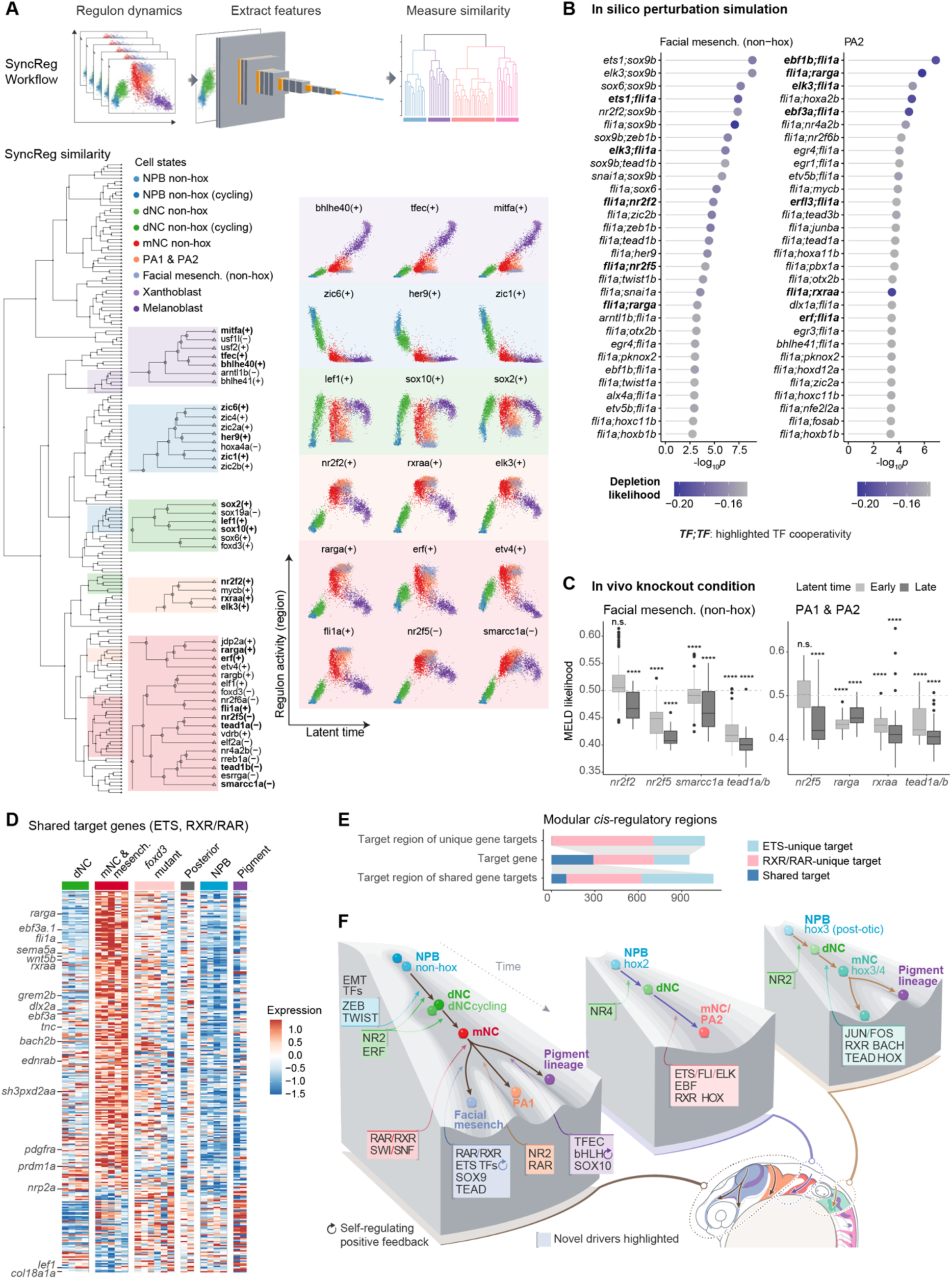
Transcriptional synergistic and redundant effects explored in regulon-mediated cell fate determination. **A**, Schematic diagram (upper panel) and results (lower panel) of SyncReg temporal dynamic analysis. SyncReg dynamics of all regulons identified temporally concurrent regulons, situated in the proximal branches of a dendrogram constructed based on dynamic similarity, with cells in dynamic trend plots colored by states. Examples of regulon temporal groups are presented, including the eight perturbed migratory/mesenchymal regulons in the orange and red panels. **B,** TF pairs of top two-gene perturbation effects ranked by the -log_10_*p*, displaying the top 30 TF pairs with the highest depletion likelihood. KO, knockout. It highlights potential cooperative TF pairs. **C,** Depletion effects of single-TF knockouts for certain lineages (subtitles). The two boxes represent the early and late phases of each lineage. MELD likelihood (y-axis) < 0.5 indicates depletion. ****, *p* < 0.0001; ***, *p* < 0.001; **, *p* < 0.01; *, *p* < 0.05; n.s., not significant, by one-sample one-sided Wilcoxon test to evaluate the lineage depletion effects. **D,** Expression profiles of targets shared between ETS and RXR/RAR TFs. Each row represents a gene, and each column represents a cell state. **E,** Distribution of targets (regions or genes, indicated by y-axis) shared between ETS and RXR/RAR TFs. **F,** Conceptual diagram of the proposed regulatory model governing cranial NC cell fate decisions.

Using SyncReg, we identified diverse regulon temporal groups associated with pigment lineage commitment (*bhlhe40, tfec, mitfa*), NPB specification (*zic1, her9, zic6*), delamination (*lef1, sox10, sox2*), migration, and mesenchymal fate commitment (Figure 7A). Within the migration and mesenchymal differentiation group, regulons such as nr2f2(+), rxraa(+), rarga(+), nr2f5(-), smarcc1a(-), tead1a(-) and tead1b(-), shared similar activity patterns with elk3(+), fli1a(+), and erf(+) (Figure 7A). Our dendrogram-comparison analysis indicated that regulatory synchronization occurs independently of motif similarity or shared gene expression profiles, emphasizing the combined role of TF binding and gene expression in defining specific NC states (Figure S7A).

To validate the convergence of TF functions, we performed double-TF perturbation simulations using RegVelo, as an orthogonal computational approach. Knockouts combining ETS TFs (e.g., *ets1, elk3, fli1a*) and RXR/RAR family or NR family TFs significantly depleted facial mesenchymal lineages, while combinations between ETS TFs and RAR, EBF, TEAD TFs caused depletion in the second pharyngeal arch lineage (Figure 7B).

Hence, we examined the individual functions of these TFs, including NR (*nr2f2, nr2f5*), RXR/RAR (*rxraa, rarga*), SWI/SNF (*smarcc1a*), TEAD (*tead1a/b*, *tead3a/b*), which may act synergistically with the ETS TFs, by analyzing in vivo knockout effects using Perturb-seq data. Knockouts of *rxraa*, *rarga*, and *tead1a;tead1b* disrupted differentiation in the first and second pharyngeal arches, while *smarcc1a, nr2f2*, and *tead1a;tead1b* knockout reduced the non-hox facial mesenchymal fate (Figure 7C). Moreover, in vivo *rxraa, nr2f5, smarcc1a,* and *erf* knockouts affected both mesenchymal and non-mesenchymal lineages, suggesting their broad roles in NC development (Figure 4D). These results confirm that NR (*nr2f2, nr2f5*), RXR/RAR (*rxraa, rarga*), SWI/SNF (*smarcc1a*), and TEAD (*tead1a/b*) TF families drive cranial mesenchymal fates, converging functionally with ETS TFs.

One key question is how the endothelial-like program promotes NC-specific migration and mesenchymal fates. We hypothesized that it is achieved by synergistic functions between the ETS TFs (e.g., *fli1a, elk3*) and RXR/RAR TFs (e.g., *rxraa, rarga*), as indicated by the Jaccard index (Figure S5A). A previous study has reported the interaction between ETS TFs and the retinoic acid (RA)-dependent regulation in activating the guanylyl cyclase/natriuretic peptide receptor-A in other systems^120^, but their role in NC development remains underexplored.

According to SCENIC+-inferred GRN, most of the shared targets between ETS factors (*elk3*, *fli1a*, *ets1*) and RXR/RAR factors (*rxraa*, *rarga*) were expressed in the migratory and mesenchymal NC, and some shared targets were involved in pathways of NC cell migration (e.g., *sema5a, tnc, sh3pxd2aa, pdgfra, nrp2a*) and cartilage development (e.g., *wnt5b, grem2b, dlx2a, ednrab, prdm1a*) (Figure 7D). Topological analysis of the *fli1a-*centered network revealed mutual interactions between *fli1a* and RXR/RAR TFs (Figure S5B). To gain insights into their synergistic mechanisms, we quantified the gene and region targets shared by ETS and RXR/RAR TFs and found that the shared target genes were primarily regulated by TF binding onto different *cis*-regulatory elements (Figure 7E). This highlights a cooperative mechanism where ETS and RXR/RAR TFs modulate NC-derived mesenchymal differentiation of cartilage derivatives, as well as migration, via modular *cis*-regulatory regions^121^.

## Discussion

Our study reconstructed cranial NC trajectories and their associated GRN with single-cell spatiotemporal resolution. Using our understanding of GRNs, we demonstrated that modulating single or multiple regulons reshapes the differentiation landscape, inducing or repressing specific fates, and these findings were validated experimentally, emphasizing the previously neglected drivers such as SWI/SNF^122,123^, BACH^124^, TEAD^125^, and EBF^26^.

While prior studies explored NC GRNs, they lacked the necessary resolution to map specific states and transitions comprehensively^11,126^. Combining in silico and in vivo knockouts, we systematically reconstituted Waddington’s epigenetic landscapes^127,128^, defining and testing the lineage- and state-specific roles of essential TFs (Figure 7F). We identified stable attractor states, intermediate unstable states (e.g., dNC, mNC), and surface characterized by monostable (delamination, migration) or bi-/multi-stable surface (differentiation) within the NC differentiation landscape. Two key mechanisms were found to underlie how GRNs shape this landscape. First, chromatin dynamics, characterized by histone modifications and chromatin accessibility, demonstrated that chromatin remodeling plays a critical role in governing state transitions during cranial NC development^122,129,130^. Furthermore, impaired chromatin remodeling has significant clinical implications^131^. Second, we identified positive autoregulation circuits, such as those involving pro-mesenchymal ETS TFs, which stabilize attractor states like facial mesenchymal progenitors. Similarly, positive autoregulation of pro-pigment bHLH TFs reinforces attractor stability^43^, consistent with established functions of TF self-regulation in prokaryotes^132^.

Our study uncovered the directive role of ETS TFs in NC migration and mesenchymal differentiation, highlighting temporal and regulatory distinctions between EMT and endothelial-like ETS-mediated migration programs. Although previous studies primarily focused on downstream cellular machinery that assists cell migration, including signaling pathways^133–135^, extracellular matrix^136,137^, and cytoskeletal rearrangement^138^, the transcriptional machinery driving this process remained overlooked. In addition, it was often oversimplified that EMT TFs drive migration^65^. This discrepancy leaves a significant gap in our understanding. By integrating single-cell multi-omics, spatial transcriptomics, in silico perturbations, and CRISPR/Cas9 knockouts, we demonstrated that EMT activation precedes migration, with post-delamination cranial NC cells adopting an endothelial-like phenotype characterized by the activation of ETS TFs such as *fli1a* and *elk3*. Such reutilization of the endothelial-like program was further supported by the activation of the FOX:ETS motif, reported to drive angiogenesis^107^ and associate with lymphomas^27^.

Our methodology efficiently discerns functional regulons even when the TFs share similar motifs, which was a major challenge in developmental studies^139,140^. Our analysis unraveled that ETS TFs regulate migration and mesenchymal differentiation through synergistic and redundant interactions. For example, *erf;erfl3* and *fli1a*;*elk3* played convergent roles in NC migration but employed distinct regulatory mechanisms. Our single-cell multi-omic approach addressed functional redundancy that previously masked ETS importance in NC migration, particularly in zebrafish models, where previous findings emphasized vascular phenotypes^112^ and did not uncover NC-specific functions. Combined with our recent findings of a toggle-switch model involving *elf1* and pro-mesenchymal ETS TFs^43^, our work extended the known functions of ETS TF interplays in embryonic development, expanding on their established contributions to angiogenesis^112^ and prostate oncogenesis^141^.

We demonstrated that ETS TFs collaborate with retinoic acid receptors (RAR/RXR) through modular *cis*-regulation, addressing gaps in understanding NC-specific combinatorial regulation, i.e., co-regulation^142,143^. We speculate a synergistic mechanism by which ETS TFs recruit CBP/p300^144,145^ to regulate acetylation, while retinoic acid receptors recruit co-factors like NCoA6/UTX/ASH2L^144^ to mediate H3K4 trimethylation (transcriptional activation) and H3K27 demethylation (repressive signal removal). Retinoic acids are important for development^76,146,147^, and previous phenotypic analyses in clinical cases and mice state that retinoic acid embryopathy results in craniofacial, cardiac, and thymic anomalies, likely through influencing cranial NC^148,149^. Our findings provide direct evidence that retinoic acid receptors mediate cellular migration and differentiation in all cranial NC lineages, primarily through synergistic effects with ETS TFs. We postulate that the NC migratory program’s control has been co-opted from endothelial cells and made more robust through redundancy and synergistic collaboration with other NC specifiers.

Our detailed view of NC migration and mesenchymal differentiation reveals important implications for congenital syndromes and NC-derived cancers. First, we established a link between ETS dysregulation in NC and craniosynostosis, as well as other NC-derived diseases. While the role of ETS TFs in craniosynostosis is well-documented^75^, their association with NC has not been well recognized. Our single-cell and whole-embryo analyses demonstrate a causal connection between the ETS TF, *ERF*, and dysfunctional migration. This highlights how disruptions in early NC development can contribute to craniosynostosis, which is typically studied at later developmental stages within facial mesenchymal cells^150^. Moreover, our study identifies *fli1a* and *elk3* TFs directly modulating NC migration, a program likely co-opted in tumorigenesis and cancer metastasis, as amplification or ectopic activation of ETS TFs, e.g., *FLI1*, is implicated in tumorigenesis of Ewing sarcoma^10,74^. The FOX:ETS motif, critical for NC migration in our study, has been strongly associated with sarcoma in previous analysis^27^. By elucidating ETS TF roles in NC migration, our work implies a co-option of endothelial-like migration mechanisms in cancer progression, with particular relevance to cancer progression^111^ and its potential as a therapeutic target^151^.

In summary, our study presents a comprehensive analysis of cranial NC developmental lineages, linking cellular states with molecular regulatory circuits and elucidating the regulatory architecture governing cell fate decisions. By integrating GRN reconstruction, spatiotemporal analysis, and regulon modulation, we provide a robust framework for understanding NC biology and highlight its significance in congenital syndromes and cancer.

## Supporting information

Table_S1_Sample information

Table_S2_Marker genes of NC cell states

Table_S3_NC GRN

Table_S4_Biological processes functional cluster

Table_S5_GOEA of BACH, EBF and TEAD regulons

Table_S6_Primer sgRNA spacer

## Acknowledgments

We thank the MRC WIMM Flow Cytometry Facility, MRC WIMM Advanced Single Cell OMICS Facility, MRC WIMM Centre for Computational Biology, MRC WIMM Sequencing Facility, and Wolfson Imaging Centre for their help and technical support in this study. Z.H. also thanks the HPC core facility of the Medical Research Institute at Wuhan University for their technical support. We thank Dr Vanessa Chong-Morrison, Dr Oana Pelea, and Dr Martyna Lukoseviciute for their help in MERSCOPE experiments and sgRNA design and we thank Dr Kate Attfield for providing the platform for some 10x runs. We thank A. Andersen for invaluable advice on the manuscript. Scientific illustrations (Figures 1a, 2a, 4a-b, 7a, and 7f) were created by Uta Mackensen. This research was supported by the Wellcome Trust ref 215615/Z/19/Z (to T.S.-S) and Stowers Institute for Medical Research Institutional Support (to T.S.-S). For the purpose of Open Access, the author has applied a CC BY public copyright license to any Author Accepted Manuscript version arising from this submission.

## Author contributions

Conceptualization: T.S.-S., Z.H.; Methodology: Z.H., T.S.-S.; Investigation: Z.H., S.M., W.W., J.M.S.-P., F.T., T.S.-S.; Visualization: Z.H., T.S.-S., W.W.; Funding acquisition: T.S.-S.; Project administration: T.S.-S.; Supervision: T.S.-S.; Writing – original draft: Z.H., T.S.-S., W.W., S.M.; Writing – review & editing: Z.H., T.S.-S.

## Methods

### Zebrafish husbandry

The handling of animals followed protocols approved by the UK Home Office, in compliance with UK law (Animals [Scientific Procedures] Act 1986) and adhering to the guidelines outlined in the Guide for the Care and Use of Laboratory Animals. All research involving vertebrate animals occurred at the Oxford University Biomedical Services facilities. Adult fish were cared for following established procedures^152^. In summary, the adult fish were maintained under the following conditions: a 14-hour light and 10-hour dark cycle (with lights on from 7 am to 9 pm and off from 9 pm to 7 am), housed in a closed recirculating water system at a temperature range of 27-28.5°C, fed 3-4 times daily, and maintained at a density of 5 fish per 1 L of water. Embryos were staged according to previously published methods^153^. Embryo staging was conducted under dissecting stereomicroscopes (Olympus).

### Embryo dissociation and cell sorting

Zebrafish embryos develop rapidly and externally, enabling the analysis of contiguous time points in NC development over hours and in thousands of isogenic animals. Embryos were obtained by crossing two zebrafish lines Gt(foxd3-mCherry)^ct110R^ and Gt(foxd3-Citrine)^ct110^ and incubated until desired stages. Embryos were dissociated with Liberase TM (Roche) for 25 min at 32°C. The cell suspension was filtered by a 100-µm mini strainer (PluriSelect) and stained with 1-3 µg/ml DAPI. Live single cells that were either Citrine-positive (Citrine+Cherry-) or double-positive (Citrine+Cherry+) were isolated using fluorescence-activated cell sorting (FACS) on an MA900 instrument (Sony).

### 10x Multiome sequencing

#### Single-cell multi-omics sequencing

To isolate nuclei, the Low Cell Input Nuclei Isolated protocol (10x Genomics, CG000365) was followed. The sorted cells were washed once with 0.04% BSA and then cells were lysed by adding 45 µl chilled lysis buffer (10 mM Tris-HCl pH 7.4, 10 mM NaCl, 3 mM MgCl_2_, 0.1% Tween-20, 0.1% IGEPAL CA-630, 0.01% Digitonin, 1% BSA, 1 mM DTT and 1 U/µl RNase inhibitor) into 50 µl cell suspension and keeping cells on ice for 20 s. Cell lysis was stopped by adding chilled wash buffer (10 mM Tris-HCl pH 7.4, 10 mM NaCl, 3 mM MgCl_2_, 0.1% Tween-20, 1% BSA, 1 mM DTT and 1 U/µl RNase inhibitor), centrifuging 500 rcf for 5 min at 4°C and resuspended in chilled diluted nuclei buffer (1x Nuclei Buffer, 1 mM DTT, 1 U/µl RNase inhibitor). The nuclei were counted, and quality checked by using Trypan Blue and C-Chip (NanoEnTek). The single-cell multiome ATAC libraries and gene expression libraries were prepared and sequenced by following the Chromium Next GEM Single Cell Multiome ATAC + Gene Expression User Guide (10x Genomics, CG000338).

#### Single-cell multiome data pre-processing

Fastq data from 10x Multiome were processed using cellranger-arc count v2.0.0 and mapped to a customized GRCz11 Ensembl 105 reference genome, which incorporated sequences of Citrine and mCherry for the specific fish genotypes. The ambient RNA was removed from each 10x sample via SoupX’s autoEstCont and adjustCounts functions^154^. In GRCz11 Ensembl 105, ENSDARG00000099849/*ebf3a* on chromosome 14 corresponds to *ebf1a* on ZFIN, and ENSDARG00000100244/*ebf3a.1* on chromosome 12 corresponds to *ebf3a* on ZFIN. To avoid confusion, we adopted the naming convention used by GRCz11 Ensembl 105.

In quality-control steps, cells were filtered based on multiome-snRNA-seq and multiome-snATAC-seq data quality. For RNA data, quality parameters were selected as nCount_RNA between 1000 to 15000, nFeature_RNA between 500 and 6000, and the percentage of mitochondrial genes (percent.mt) below 20%^155^. Following these quality checks, RNA-based doublets were identified and removed using the R package DoubletFinder^156^.

The multiome-snATAC-seq data underwent pre-processing using the R package ArchR^157^. The bespoke ArchR reference genome was created by incorporating BSgenome.Drerio.USCS.danRer11, the custom-modified GRCz11 Ensembl 105 reference genome, and the DANIO-CODE genome blacklist^158^. The blacklist was adapted from DanRer10 via the UCSC Liftover tool (https://genome.ucsc.edu/cgi-bin/hgLiftOver). ArchR arrow files were generated from the cellranger-arc output named atac_fragment.tsv.gz, applying a minimum transcription start site enrichment score (minTSS) of 4 and a threshold of 2500 for the minimum number of mapped ATAC-seq fragments per cell (minFrags). Multiome-ATAC-based doublets were detected using ArchR’s addDoubletScores function, using parameters of k = 10 and the LSI “tf-logidf” method, and filtered out by the filterDoublets function. Cells meeting the criteria for both RNA and ATAC quality control, as well as passing doublet filtering, were selected for downstream analysis.

### 10x Multiome genotyping

#### Multiome genotyping amplicon-seq

To overcome the issue of limited cell numbers at the early stage, we multiplexed Citrine-positive and double-positive cells for one 10x run. Although only s8 (75% epiboly - 3 ss) contained mixed genotypes, other single-genotype samples (s1 - s7) underwent the same amplicon-seq procedure to be used as the training sets. From the mixed-genotype and other single-genotype samples, amplicon-seq libraries were prepared from the 10x cDNA samples using NEBNext Ultra II Q5 Master Mix (NEB) and 10µM primers (IDT). This panel contained 7 amplicons, covering Cherry (n = 2), *foxd3* mutated allele (n = 2) and *foxd3* wild-type allele (n = 3). The PCR program was 98°C for 30 s, 25 cycles of (98°C for 10 s, 68°C for 30 s, 72°C for 30 s), 72°C for 2 min and hold at 10°C. The PCR products were purified using AMPure XP beads (Beckman Coulter). The amplicon-seq libraries were sequenced by the MiSeq 600-cycle platform (Illumina).

#### Genotype demultiplexing of 10x Multiome

Our amplicon-seq samples comprised a training set (s1 - s7) with known genotypes and a testing set (s8) for which genotypes were unknown. To create the reference genome for genotyping, we assembled the wild-type and mutant *foxd3* allele reference sequences by applying Trinity^159^ on bulk RNA-seq data GSE106676^49^. Then the amplicon-seq fastq data were aligned to these reference sequences and quantified using the Alevin function of Salmon v1.8.0^160^. Salmon output was organized into a count matrix for NC cells in training or testing sets respectively, where the rows were amplicons, and the columns were cells. The count matrices were transformed by log_2_(x+1) transformation.

To construct the model to classify genotypes, we fitted a logistic regression model using the training set with the glm function and parameters of family = “binomial” or “poisson”, as *g_i_* ∼ β_0_ + β_1_β_1,*i*_ + ⋯ + β_*n*_*e_n,i_*, where *g_i_* indicated the genotype (wild type or mutant) of the *i*-th cell, and *e_k,i_* indicated the expression level *log*_2_(*count* + 1) of *k*-th amplicon in the *i*-th cell. The model performance was evaluated using the roc function from pROC package^161^. The binomial model (AUC: 0.781, 0.742-0.820) slightly outperformed the Poisson model (AUC: 0.764, 0.724-0.804). The best cutoff of the binomial model was then decided by using the ROCR function^162^.

Next, this logistic classification model was applied to the testing set and classified the cells into the wild-type (“Citrine-positive”) or the mutant (“double-positive”) groups. For the cells that were not captured in the amplicon-seq data, we assumed that cells with the same genotype tend to share similar expression profiles. Thus, the *k*-nearest-neighbour method (function class::knn) was used to predict missing values based on the Seurat-calculated PCA space (Figure S1D).

### Spatial transcriptomic analysis

#### MERSCOPE sample preparation and experiments

Zebrafish embryos were collected, dechorionated, and deyolked before being embedded in the optimal cutting temperature (OCT) compound (VWR). OCT blocks were snap-frozen on the dry ice/ethanol bath and sectioned at 10-µm thickness onto MERSCOPE slides using a microtome-cryostat (Leica). Sections were hot fixed with 4% PFA at 37°C for 30 min, following the MERSCOPE Fresh and Fixed Frozen Tissue Sample Preparation User Guide, and imaged with the MERSCOPE Instrument (Vizgen).

#### MERSCOPE gene panel design

To identify cell types and NC cell states using MERSCOPE, we designed a gene panel comprising both global markers for non-NC cells and specific marker genes for NC cell states. We used SPAPROS, a probe selection algorithm, on multiome-snRNA-seq data, Smart-seq3 data and the organogenesis dataset of 10-, 14-, and 18-hpf embryos^29,163^. The reads per kilobase million (RPKM) and transcripts per million (TPM) count matrices were computed for stage-specific pseudo-bulk samples to filter genes based on expression levels. After submitting the initial 304-gene panel to Vizgen for a designability check, redundant markers for the same non-NC cell type were identified and eliminated, resulting in a refined set of 300 genes. These selected gene sets underwent validation using SPAPROS metrics and were assessed for their ability to delineate clusters in low-dimensional representation.

#### Assigning cell type identities

Spatial transcriptomic data were visualized using MERSCOPE Visualizer software (Vizgen) for quality assessment. Suboptimal segmentation results were addressed by re-segmenting based on polyT and DAPI using VPT and Cellpose2 human-in-the-loop methods^164^. Re-segmented data were exported to h5ad format using Visualizer and preprocessed with Squidpy^165^, filtering cells with minimal counts of 10 and minimal volume of 50. Clustering was performed using the Leiden algorithm with the first 20 principal components and a resolution of 1.5. Major cell types were identified by mapping organogenesis data^29^ onto spatial transcriptomic data with Tangram^40^. NC-dominating clusters were isolated and re-clustered with a resolution of 1.5 and 5-nn graph. The Tangram function map_cells_to_space was employed with uniform density_prior in clusters mode. To distinguish NC-derived from non-NC-derived head mesenchymal cells, we used NC markers, namely *grem2b, foxd3, sox10, tfap2a, zeb2a*, and *crestin*. Cells expressing at least one NC marker were categorized as NC derivatives, while those lacking NC marker expression were assigned to non-NC-derived head mesenchyme. Cell locations were visualized using Squidpy’s spatial_scatter function. We termed broad NC states into NC (mesenchymal/mesench.), i.e., predominantly grem2-high NC^29^, and NC (other), i.e., predominantly crestin-high NC^29^. Neighbor enrichment in wild-type and knockout embryos was evaluated using the gr.spatial_neighbors and gr.nhood_enrichment functions from the Python module Squidpy^165^.

### ChIP-seq and data analysis

#### MicroChIP-seq

To validate the SCENIC+-predicted CREs, the related histone modifications were profiled in wild-type and *foxd3*-mutated cells from 12-16 ss embryos. Fish lines were crossed, and embryos were incubated in the same way as the 10x Multiome samples. The manufacturer’s protocols of True MicroChIP-seq kit (Diagenode) and Chromatin EasyShear Kit - High SDS (Diagenode) and the Fish’n ChIPs (Chromatin Immunoprecipitation in the Zebrafish Embryo) protocol^166^ were followed. In brief, dissociated cells were fixed using 1% paraformaldehyde for 8 min at room temperature and stopped by glycine before FACS. Citrine-positive (wild-type) and double-positive (*foxd3*-mutated) post-fixation cells from 12-16 ss embryos were isolated by FACS. The chromatin was sheared by 3 cycles of 30-s ON/30-s OFF on the Bioruptor Pico (Diagenode). The antibodies used were H3K27me3 (Millipore, cat no. 07-449), H3K27ac (Abcam, cat no. ab4729), H3K4me1 (Diagenode, cat no. pAb037050 / C15410037), and H3K4me3 (Diagenode, cat no. pAb003050 / C15410003). Controls of ’IgG-cit’, ’IgG-dp’, or ’input-cit’ were performed.

#### ChIP-seq data analysis

Raw reads were quality-controlled using FastQC (https://www.bioinformatics.babraham.ac.uk/projects/fastqc/). Following quality control, the reads were trimmed using Trim Galore (https://github.com/FelixKrueger/TrimGalore), aligned with Bowtie2^167^, and the resulting BAM files were indexed using Samtools^168^. For peak calling and bigwig generation, each experimental condition was compared to a corresponding control, which could be ’IgG-cit’, ’IgG-dp’, or ’input-cit’. Peaks were identified from these sorted BAM files using MACS2^169^ and then annotated with HOMER^170^. Finally, Deeptools^171^ was employed to ensure data quality, normalize the data, and provide visual representations of the results. The samples that were outliers in the Spearman correlation analysis were excluded from the downstream analysis.

### RNA in situ hybridization and immunocytochemistry

#### Hybridization chain reaction (HCR)

To validate the novel NC markers identified in this study, fluorescently labeled hairpins, DNA probe sets, and HCR buffers, whether procured commercially (Molecular Instruments) or produced in-house, were used for the multiplexed targeted visualization of mRNA in situ, employing the HCR kit (version 3, Molecular Instruments)^172^. Zebrafish embryos, harvested at the desired developmental stages, underwent dechorionation and fixation in 4% paraformaldehyde at 4°C for 24 hours. The HCR procedure adhered to the ’In situ HCR v3.0 protocol for whole-mount zebrafish larvae’ (Molecular Instruments). Post HCR, the embryos were embedded in 1% low melting-point (LMP) agarose and imaged using the 780 upright, 780 LSM inverted or 980 LSM inverted confocal microscope (Zeiss). Image analysis was performed using Fiji and Zeiss Zen.

#### Whole-mount immunofluorescent staining

Embryos at desired stages were fixed with 4% paraformaldehyde (Methanol-free, Thermo Fisher) for 1 hour at room temperature, followed by rinsing in PBT (2% DMSO, 0.5% Triton-X in PBS). Blocking was carried out in a solution of 10% goat or donkey serum in PBT for one hour at RT. The primary antibodies were diluted in blocking solution: goat anti-tfap2a antibody 1:50 (LS–C87212, LSBio), rabbit anti-Elavl3/4 antibody 1:200 (GTX128365, GeneTex), and chicken anti-GFP 1:200 (ab13970, Abcam), and applied to the embryos for overnight incubation at 4°C. After thorough washes with PBT, the secondary antibodies, donkey anti-rabbit 647 nm (A31573, Thermo Fisher), donkey anti-goat 568 nm (A11057, Thermo Fisher), donkey anti-chicken 488 nm (A78948, Thermo Fisher), and Hoechst 33258 (ab228550, Abcam, 1:1000 dilution) were prepared in PBT and added to the embryos for a 2-hour incubation at room temperature in the dark. Post extensive washing with PBT, embryos were embedded for confocal imaging.

### Single-cell multiomic computational analysis

#### Clustering analyses and cell type annotations

Our first step was the pre-processing and clustering analysis of the 10x Multiome-snRNA-seq dataset using Seurat v4.2 package^155^. RNA data normalization was achieved with the “LogNormalize” method with a scale factor set at 10,000. We then selected the top 2000 highly variable features using the “vst” method. This was followed by data matrix centering, scaling, and principal component (PC) computation. For clarity, the first 25 PCs were chosen based on the elbow plot method for further analysis. Clusters were delineated using the functions FindNeighbors and FindClusters set at a resolution of 0.6. The marker genes were identified using the specific parameters (only.pos = TRUE, min.pct = 0.25, and logfc.threshold = 0.25).

For the multiome-ATAC dataset, the peak-cell count matrix was refined through dimensionality reduction using the ArchR Iterative Latent Semantic Indexing (LSI) method, followed by visualization with Uniform Manifold Approximation and Projection (UMAP). Marker peaks were identified using the ArchR getMarkerFeatures function with the Wilcoxon test method. The motif enrichment analysis of these marker peaks was conducted using the ArchR peakAnnoEnrichment function, comparing against the DANIO-CODE motifs or tailor-made motif collections.

To annotate the cell clusters, we transferred the labels from a zebrafish organogenesis scRNA-Seq dataset^29^. This involved normalizing, merging the top 2000 variable feature lists, scaling data, running PCA and computing anchors with the first 50 PCs. The labels of cluster names and tissue names were transferred by Seurat’s TransferData from the organogenesis dataset to our multiome-snRNA-seq dataset. The non-NC clusters were annotated by the dominating labels. Three clusters with low numbers of features detected were annotated as “mutant low-feature”, “neural crest low-feature” and unclassified. The neural crest clusters were identified based on the label transferring results and manually annotated as “neural crest”, “neural crest mutant”, “neural crest early”, “neural crest pigment”, and “neural crest low-feature”.

To identify potential intermediate cells between the NC and neural plate populations, we subset the multiome-snRNA-seq data for both NC and neural clusters and performed the clustering analysis. This analysis did not reveal any intermediate population, but *jhy*-expressing apoptotic clusters were identified and subsequently removed.

To retain only high-quality NC cells for further analysis, we excluded the cells expressing genes from the panel (*jhy*, *cyt1*, *krt5*, and cyt11) commonly associated with apoptosis and epidermal populations. The NC multiome-snRNA-seq dataset underwent the same pre-processing steps as the complete multiome-snRNA-seq dataset. We employed the top 15 PCs for clustering and visualization. Louvain clustering was implemented using a resolution of 1. Clusters were subsequently annotated, drawing on stages, genotypes, and cell cycle states as determined by the Seurat CellCycleScoring function. Some gene expression figures were made with the R package scCustomize (https://samuel-marsh.github.io/scCustomize/index.html).

#### Integration analysis of NC data

To compare the cell states across three datasets (zebrafish organogenesis^29^, our multiome data, and Smart-seq3 data^43^ of cranial NC cells containing seven stages from 3 ss to 22 ss and two genotypes, i.e., wild type and *foxd3* mutant), Seurat canonical correlation analysis (CCA) was used to identify anchors from the first 50 PCs for integration. Because cell populations between multiome-snRNA-seq and Smart-seq3 datasets are largely overlapped, Seurat-CCA in this scenario was unlikely affected by the overcorrection error.

To quantify the inter-group similarity, we applied the getIDEr function from the CIDER R package^173^. The organogenesis cell labels were simplified by merging clusters containing the differentiating neurons, epidermal, pharyngeal arch, diencephalon, pronephric duct, mesoderm, heart, muscle, and lateral line, respectively. The apoptotic clusters were from the organogenesis dataset. The curated annotations were used as the initial cluster and the inter-project batch effects were removed using the limma trend method^174^. The inter-group similarity matrix was visualized using the R package pheatmap (https://cran.r-project.org/web/packages/pheatmap/index.html).

#### RNA velocity analysis

For 10x Multiome data, bam files generated by cellranger-arc were processed by the velocyto run10x command to generate loom files^52^. To account for the loss of cytoplasmic mRNA and incomplete-length transcript information in the multiome-snRNA-seq data, we reproduced the multiome-snRNA-seq trajectories using the Smart-seq3 dataset. For Smart-seq3 data, the bam files for velocyto were created by zUMIs and processed by “veloctyo run --vv --umi-extension Gene -d 1 --without-umi” to use both internal reads and UMI reads.

To perform velocity analysis for wild-type NC development, the NC multiome data were divided into the wild-type and the mutant datasets. The wild-type NC data were pre-processed by the scVelo preprocessing functions filter_genes, normalize_per_cell, filter_genes_dispersion, and log1p^51^. Neighbors were computed by scVelo preprocessing functions neighbors and moments functions with n_pcs of 30 and n_neighbors of 10. RNA velocity was then estimated by the velocity and velocity_graph functions. We computed the PAGA velocity graph using scVelo with minimum_spanning_tree = False and threshold_root_end_prior = 0.5. The mutant NC data were pre-processed in the same way. To visualize the PAGA results, we used the threshold of 0.04.

#### Multiomic velocity analysis

To use both gene expression and chromatin accessibility to estimate velocity, we used MultiVelo^35^. The multiome dataset was a subset to retain only the most anterior lineage. Cells with counts between 1000 and 20000 were kept for MultiVelo analysis. The RNA data were processed as recommended by scVelo functions. As for the ATAC data, the peak-count matrix called by SCENIC+ pycisTopic was used. The feature linkage bedpe file was generated by Signac^175^. The peaks were annotated by Homer v20201202 annotatePeaks.pl and parsed by an in-house script. The ATAC data, the peak annotation file and the linkage file were used as input to the MultiVelo aggregate_peaks_10x function. Cells were filtered for ATAC counts between 1000 and 60000. The aggregated peaks were normalized by TF-IDF. RNA data were normalized again after filtering. To smooth the ATAC data, Seurat WNN of 30 *k*-nearest neighbors was used.

RNA and ATAC datasets were then used to fit the multi-omic dynamical model by function recover_dynamics_chrom with max_iter=5 and init_mode=“invert”. The latent time was computed using MultiVelo velocity_graph and latent_time functions. The pie summary plot was visualized by the MultiVelo pie_summary function. Thus, 30 nearest neighbors were used.

#### Lineage inference and probability calculation

To compute the fate probability, MultiVelo velocity vectors were used as input of Cellrank2^104,105^ to quantify the transition matrix with the ‘deterministic’ model. To refine the clusters, we applied Leiden clustering using a resolution of 1 and manually curated and merged the clusters. Next, the estimator was initialized and fit using the Schur decomposition and GPCCA algorithms to estimate macrostates by 8 states. The terminal states were manually set to ‘Pigment_sox6_high’, ’mNC_arch1’, ’mNC_head_mesenchymal’, ’Pigment_gch2_high’, and the initial state was predicted to be “NPB nohox 1”. Next, the fate probability matrix was computed, and the probabilities were extracted from the “obsm[’terminal_states_memberships’]” entry. For visualization, we used cellrank.pl.heatmap and cellrank.pl.gene_trends.

#### De novo motif discovery for the *foxd3* motif

Because the *foxd3* motif in the Danio Code motif database failed cisTarget analysis, we reasoned that the ortholog-based motif was not applicable in the NC development scenario. Thus, we sought to *de novo* construct the *foxd3* motif from the biotin-ChIP data ^176^. The *foxd3* biotin-ChIP raw data (SRR6268024/5/6/7 as ChIP sample and SRR6268028/29/30/31 as input control) were downloaded from the Sequence Read Archive (SRA). Raw data were preprocessed by Trim Galore v0.6.5 and aligned by Bowtie2 v2.4.2 to GRCz11. The duplicates were removed by Samblaster v0.1.24^177^. The narrow peaks were called from the biotin-ChIP sample and the input control using MACS2 and the regions in the blacklist were excluded. Subsequently, we used two tools, GLAM2 from MEME v5.4.1 and MEMEChIP, to predict motifs from the MACS2-called peaks^178,179^. Based on the hypothesis that the *foxd3* motif should be enriched in the migratory NC populations, we identified a GLAM2 motif (glam_motif1), which satisfied the enrichment and resembled the orthologous FOXD3 motif “5′-TGTTTGTTT-3′”. This GLAM2*-*predicted *foxd3* motif was used for the SCENIC+ analysis.

### GRN analysis by SCENIC+

The comprehensive GRN underlying both normal and *foxd3*-mutant development was constructed using SCENIC+^33^. SCENIC+ could integrate both snRNA-seq data and snATAC-seq data to infer enhancer-driven regulons.

Our approach involved preparing scRNA-seq data, the snATAC-seq pycisTopic object, and the pycisTarget motif enrichment dictionary to create the SCENIC+ object. Our dataset contains 10,604 wild-type and 5,946 mutant NC cells. For the multiome-snRNA-seq dataset, preprocessing involved adjusting the *foxd3* read counts to 10% in the *foxd3*-mutated cells to emulate the loss of function. The multiome-ATAC dataset underwent preprocessing with pycisTopic, wherein CellRanger-ARC fragment output files were processed through a series of pycisTopic functions (export_pseudobulk, peak_calling, get_consensus_peaks, compute_qc_stats, and create_cistopic_object_from_fragments) to form the pycisTopic object. To determine the optimal number of topics in the topic modeling, the parameter sweeping was done from 5 to 100 in increments of 5, with the 95-topic model ultimately being chosen. Post topic modeling, differentially accessible regions per cell type, serving as enhancer candidates, were identified from the pycisTopic object.

Subsequently, the cisTarget database was established for pycisTarget analysis. Because the DANIO-CODE study focused on the conservation of DNA binding regions over general ortholog information, the motif database incorporating DANIO-CODE motifs along with the de novo *foxd3* motif was selected for pycisTarget analysis. The feather database was constructed using scripts from the create_cisTarget_databases repository. Motif enrichment analysis on the enhancer candidates predicted by pycisTopic was conducted using the SCENIC+ wrapper function run_pycistarget.

The SCENIC+ object was created by merging three key components: RNA data, the pycisTopic object, and the motif enrichment results. To reconstruct the GRN, the step-by-step SCENIC+ protocol was followed, including the steps of cistrome generation, region-to-gene and TF-to-gene relationship calculations, and actual GRN construction. To expedite the region-to-gene and TF-to-gene relationship calculations, a stochastic gradient boosting model (sGBM) was employed, configured with a learning rate of 0.01, 5,000 estimators, max features set at 0.1, and subsample rates of 0.5 and 0.2 for region-to-gene and TF-to-gene calculations respectively. The observation subsampling fraction was reduced to 0.2, acknowledging the relatively large number of observations in our dataset.

Following GRN construction, the activity of gene-based and region-based eRegulons (AUCell scores) was quantified using the function score_eRegulons, and their regulon specificity scores (RSSs) were derived using the function regulon_specificity_scores. The network was exported by SCENIC+ functions (create_nx_graph and export_to_cytoscape) and visualized by Cytospace (https://cytoscape.org). To check the overlap between regulons, the Jaccard indexes were calculated using SCENIC+ jaccard_heatmap function as *J*(*R_1_*, *R_2_*) = |*R*_1_ ∩ *R*_2_|⁄|*R*_1_ ∪ *R*_2_|, where *R*_1_ and *R*_2_ represent the sets of elements in regulon 1 and regulon 2, respectively.

### GRN-informed velocity analysis by RegVelo

RegVelo was a novel velocity model that combined GRN with the RNA velocity framework by modeling the transcription rate of each gene as governed by upstream TFs. It simulated the full dynamics from gene transcription to RNA splicing and learned the entire process through deep generative neural networks.

To study the underlying connection between cell differentiation dynamics and gene regulation, we used the SCENIC+-inferred GRN to assist RNA velocity inference through RegVelo. In the RNA velocity framework, the dynamic process of splicing for each gene was modeled using the chemical master equation (equation 1):

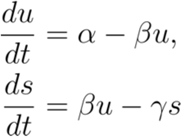

Here, *u* and *s* represented size-normalized abundances of unspliced and spliced counts, respectively. The model parameters α, β, and γ corresponded to the RNA transcription, splicing, and degradation rates. Traditionally, α was assumed to be a constant^52^ or binary state (ON or OFF)^51^. In RegVelo, the coefficient α was modeled according to the following equation (equation 2):

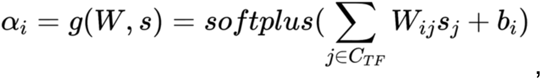

and *b_i_* represented the base transcription level.

Here, *C_TF_* represented the sets of all TFs, and *W* represented the regulation coefficient (GRN) learned by RegVelo, where columns corresponded to the regulators and rows corresponded to the targets. This equation allowed us to incorporate gene regulation learned from SCENIC+ into RNA velocity, assuming that those target genes would be regulated by upstream TFs if the regulatory relationship was present in the SCENIC+ GRN. Additionally, RegVelo learned the gene-specific latent time, and integrated the latent time with velocity equation (1) to get the predicted spliced and unspliced readout. All kinetic parameters and latent time were learned through the variational autoencoder.

RegVelo was applied to both 10x Multiome and Smart-seq3 datasets. For the Smart-seq3 dataset, we used the “hard constraint” mode, which assumed that each target gene transcription rate was fully regulated by the SCENIC+-inferred upstream regulators. For the 10x Multiome dataset, RegVelo was applied to the non-hox region dataset. Given that our GRN was learned from the full NC dataset, we aimed to prevent missing region-specific gene regulation. Therefore, we used the “soft constraint” mode in RegVelo, which learned new regulatory connections in addition to the prior GRN.

### Tn5-bias-corrected de novo motif discovery

To aid de novo motif discovery in our multiome-ATAC data, we used ChromBPNet to mitigate Tn5 bias. ChromBPNet used convolutional neural networks to predict base-resolution cluster-specific chromatin accessibility profiles, enabling precise in vivo motif identification^68^. Key cell states in cranial NC trajectories were represented by selecting clusters including merged_dNC_nohox, merged_NPB_nohox, mNC_nohox, mNC_head_mesenchymal, mNC_arch1, Mutant_nohox_early, Mutant_nohox_late, and Pigment_sox6_high. To secure sufficient depth for dependable Chrombpnet model training, these clusters comprised cell counts exceeding 950, except for Mutant_nohox_late (N = 663) and Mutant_nohox_early (N = 828).

Pseudo-bulk bam files were generated from cellranger-arc bam and cluster annotations using pyflow-scATACseq (https://github.com/crazyhottommy/pyflow-ATACseq). MACS2 was employed for peak calling with a significance threshold of q = 0.01 and an effective genome size of 1.4e9, and the identified peaks were systematically filtered using the blacklist. Chromosomes were divided using the “chrombpnet prep splits” command with the Fold 0 parameter, defining training chromosomes (chr1, chr3, and chr6) and validation chromosomes (chr8 and chr20). Non-peak background regions were created by “chrombpnet prep nonpeaks”. Cluster-specific ChromBPNet Tn5 bias models were trained using ’chrombpnet bias pipeline - d “ATAC” -b 0.5’, followed by training cluster-specific bias-factorized models through ’chrombpnet pipeline -d “ATAC”’ with cluster-specific Tn5 bias models. Contribution scores were predicted by “chrombpnet contribs_bw” with the bias-factorized models.

After contribution score prediction, de novo motif discovery was conducted using the TFmodisco-lite command ’modisco motifs’ with the input of predicted contribution profile scores^106^. All newly discovered motifs were concatenated and clustered using the Gimme command ’gimme cluster -t 0.999 -N 4’^180^. For motif annotation, the average motifs were meticulously compared against the DANIO-CODE motifs via TOMTOM ’tomtom -no-ssc -oc . -verbosity 4 -text -min-overlap 5 -dist pearson -evalue -thresh 10.0’^181^, followed by manual curation to eliminate artificial motifs and reduce redundancy. Compared to the publicly available motif databases, the de novo discovered motifs might better represent in vivo scenarios. However, the cluster-specific discreet pseudo-bulk samples used by motif prediction can undermine the variability along continuous trajectories. Thus, de novo motif analysis and SCENIC+ GRN reconstruction were applied to be complementary to each other. The hit calling of the de novo TFmodisco-lite motifs was conducted using Finemo^68^. The motif enrichment was performed using the converted position weight matrices (PWMs) in R package ArchR^157^.

### Functional and temporally dynamic analysis of regulome

#### Functional enrichment analysis

To systematically analyze regulon functionality, SCENIC+-predicted regulons were initially grouped into broad functional clusters through agglomerative clustering, utilizing the regulon-based AUCell matrix (Figure S3D). For an in-depth interpretation of these regulon clusters, FishEnrichR^182^ was employed to conduct gene ontology enrichment analysis on the target genes within each regulon cluster. This approach helped identify regulon clusters that played potential roles in early NC, pigment lineage, NC migration, hindbrain NC, cardiac NC, posterior NC, and chromatin remodeling.

#### Centrality analysis

Within each cluster, the TFs of the regulons were prioritized based on the out-degree centrality of the TF-TF sub-network, which was calculated by applying networkx.out_degree_centrality function on the TF-only skeleton of the SCENIC+-predicted GRN^183^. High out-degree centrality in the TF-TF network indicated higher positions of TFs in the GRN hierarchy (Figure 3C); however, this method did not incorporate temporal information.

#### Motif similarity analysis

To assess motif similarity, we first generated the Markov background model (order 3) using MEME (version 5.4.1) with the command ‘fasta-get-markov -m 3’. Next, we converted the position weight matrix (PWM) format to MEME format using the chen2meme function. Motif similarity was computed using TOMTOM with the parameters ‘-dist kullback -motif-pseudo 0.1 -text -min-overlap 1’^181^.

#### Temporally dynamic analysis of GRNs by SyncReg

To capture the dynamic nature of NC development, we designed an integrative analysis combining temporal metrics and regulon activities, termed “regulatory synchronization (SyncReg)”. This approach, akin to temporal gene trend analysis, offered the advantage of minimizing the influence of dropout in regulon-level assessments.

Initially, we evaluated three algorithms, scVelo^51^, MultiVelo^35^, and RegVelo, to determine the best performer in predicting latent time for non-hox trajectories. This comparison involved two steps. Firstly, we correlated the predicted latent time with the ordinal variable representing development stages (“real-world” time) using Spearman correlation coefficients. Next, we compared the predicted latent time between pairs of methods to discern differences in their predictions. Lastly, we selected MultiVelo over scVelo due to its higher Spearman correlation coefficient and better representation of dNC-mNC transition (Figure S6A-C). Although MultiVelo and RegVelo showed similar performance in latent time prediction, we opted for MultiVelo to capture chromatin changes over time.

We integrated MultiVelo-predicted latent time (x-axis) with regulon activities assessed by AUCell scores (y-axis) into a two-dimensional framework. Feature extraction from this dynamic framework was conducted using a VGG16 model^184^, followed by cosine similarity calculation between each pair of regulons. Subsequently, hierarchical clustering with the complete-linkage method was applied to group the regulons based on the resulting similarity matrix. Dendextend visualization facilitated the comparison of different hierarchical clustering outcomes, providing a refined understanding of regulon interactions and temporal behavior.

The temporal activity of individual regulons was visualized using the R package ggplot2, with trend estimation SCENIC+ AUCell methods quantified the regulon activities, and cluster-specific regulons were identified based on the top twenty regulons for each cluster, determined by gene-based RSSs.

### In silico perturbation simulation

Traditional perturbation simulation methods like CellOracle^34^, Dynamo^185^, and SCENIC+^33^ typically begin by constructing a predictive function for target gene expression (or velocity) *f*(*TF*_1_, *TF*_2_, …, *TF*_*k*_) = *f*(*TF*_1_) + *f*(*TF*_2_) + ⋯ + *f*(*TF*_*k*_), using regulator gene expression as input. They then reduced the expression level of the TFs of interest and propagated the perturbation effects to other genes through this function *f*(*gene expression*).

In contrast, RegVelo predicted target gene expression by integrating RNA kinetics over latent time, enabling the prediction of long-term effects and aligning more closely with TF knock-out experiments in vivo.

Assuming we had a fitted RegVelo model, the gene velocity was defined as the following coupled model:

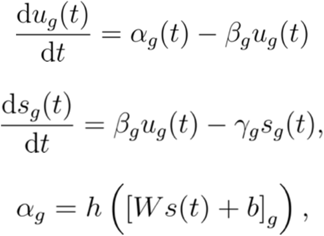

Unlike other methods, RegVelo generates a perturbed vector field de novo by introducing GRN perturbation. Specifically, we can mask the regulon (downstream targets) of each TF. For example, if we wanted to create perturbation to *TF_k_*, we created a perturbed weight matrix *Ŵ* as follow:

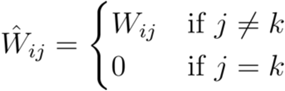

We then replaced the original weight matrix with the perturbed weight matrix, performed integration, and generated a new velocity vector field *v̂*(*t*). Based on these two vector fields, we used CellRank2 to predict fate probability towards the terminal state. Consequently, we obtained two fate probability matrices Π ∈ ℝ^*n×p*^, Π̂ ∈ ℝ^*n×p*^, where each row represented a cell, and each column represented the terminal state. Finally, we could define the *deletion likelihood* ℓ_*d*_ of *TF*_*k*_ to the *m*-th terminal state. This statistic quantifies the probability that *m*-th terminal state will be depleted after *TF*_*k*_ knockout. To compare depletion effects across different lineage, RegVelo introduces a rescaled score named *depletion score* to indicate the perturbation effect:

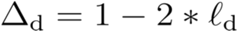

This score indicated the perturbation effects when we masked the *TF_k_* regulon to the cell dynamics. If Δ_*d*_ > 0, it indicated enrichment. If Δ_*d*_, < 0, it indicated depletion.

Compared with previous perturbation methods, RegVelo offered several advantages. First, it modeled gene expression from full RNA kinetics rather than simple gene expression regression, allowing for the simulation of long-term effects. Second, unlike methods like CellOracle that calculated perturbation effects in 2-dimensional space, RegVelo generated perturbed single-cell dynamics at the gene level. By combining this with CellRank2 to simulate deviations in fate probability within each cell, RegVelo achieved finer-grained and more reliable quantification of perturbation effects. Lastly, Unlike Dynamo, where gene regulation inference was solely based on estimated RNA velocity without prior GRN input, RegVelo incorporated prior knowledge, resulting in a modeled regulatory system with higher confidence and more accurate modeling of perturbation effects.

### In vivo Perturb-seq

#### Design of chemically modified sgRNA

The sgRNA designs were sourced from various origins, including CHOPCHOP in-house design^186^, the IDT sgRNA design tool, and existing literature. Specifically, the sgRNAs for *fli1a*, *tfec*, *rxraa*-sgRNA6, *rarga*-sgRNA3, and *erfl3*-sgRNA1 were derived from the IDT sgRNA design tool. The *nr2f5*-sgRNA4 corresponded to ZDB-CRISPR-210310-12^187^, while the *tyr* sgRNAs were obtained from the MIC-Drop study^36^. The selection criteria for sgRNAs, adapted from the same study^36^, included: a) 2-4 sgRNAs were employed per gene to maximize overall efficiency; b) spacer sequences were fixed at 20 nucleotides (or 19 if 20-nt spacers were unavailable); c) spacer sequences were required to have no fewer than three mismatches in off-target alignments; d) in cases where spacer sequences had off-target alignments with three mismatches, these mismatches were mandated to occur in the seed region for designs executed using CHOPCHOP. Highly expressed onhologs^188^ were also considered for multi-gene knockouts. Details of the spacer sequences for sgRNAs are cataloged in Supplementary Table 6.

As the ability to be template switched is the key to sgRNA capture rates, three different designs of sgRNA were tested for their template switching efficiency (Figure S4A, B). There were no obvious differences. The Alt-R sgRNA design was chosen due to cost efficiency and durability in cells.

#### Ribonucleoprotein microinjection and generation of F_0_ Crispants

The ribonucleoprotein (RNP) mixture was prepared by incubating the combination of sgRNAs, EnGen Cas9 (NEB), and nuclease-free water at room temperature for 10 min, followed by cooling on ice. Phenol red was added to the RNP mixture prior to injection. The final injection mixture contained 250 ng/µl sgRNAs (approximately 7.8 µM), 3.36 µM EnGen Cas9, and 0.05% (w/v) Phenol red. Embryos obtained by crossing Gt(*foxd3*-mCherry)^ct110R^ or Gt(*foxd3*-Citrine)^ct110^ and wild-type fish were injected with 1-2 nl of the injection mixture and incubated until reaching 20-21 ss.

#### Validation of sgRNA knockout efficiency

To assess the effectiveness of RNP injection, embryos at the 1-cell stage were injected with RNP containing three mCherry sgRNAs. The presence of fluorescence in the embryos was then examined one day post-injection using the MVX10 stereomicroscope (Olympus) (Figure S4C). A complete elimination of mCherry fluorescence indicated successful knockout.

Furthermore, the knockout efficiency was also evaluated using RNP injections targeting *tyr* (Figure S4D). Embryos at a certain post-injection day were embedded in low-melting-point agarose and subjected to imaging with the MVX10 stereomicroscope (Olympus) to confirm the efficacy of the gene disruption.

#### Direct capture Perturb-seq

For the Perturb-seq process, embryo dissociation was performed similarly to the 10x Multiome. Citrine-positive live cells were sorted into 100 µl of 0.1% UltraPure BSA (Invitrogen) in PBS and then centrifuged at 600× g for 5 min at 4°C. Care was taken to remove the supernatant without disturbing the cell pellet, leaving approximately 40 µl of supernatant. The cell pellet was reconstituted in the remaining supernatant. Cell count and quality were assessed using Trypan Blue and C-Chips (NanoEnTek). The protocol CG000510 Chromium NextGEM Single Cell 5’ v2 CRISPR User Guide (Rev B, 10x Genomics) was adhered to for single-cell gene expression and CRISPR library preparation. Sequencing of the libraries was carried out on NextSeq and NovaSeq platforms (Illumina).

#### Perturb-seq data pre-processing

The raw datasets underwent preprocessing with the ’cellranger count’ function from CellRanger (version 7.1.0), using the SC5P-R2 chemistry. Sequencing reads were aligned to the custom reference genome, which integrated the GRCz11 Ensembl 105 reference with additional entries for the mCherry and Citrine genes. Cells were demultiplexed into distinct knockout panels based on sgRNA counts over one. Due to the specific arrangement of panels and 10x sequencing runs in our study, each 10x run was devoid of nested knockout panels. Consequently, cells harboring sgRNAs from two disparate knockout panels were identified as doublets. Ambient RNA contamination was regressed on a sample-by-sample basis using SoupX^154^. RNA-based doublet cells were identified using DoubletFinder^156^, and doublets and sgRNA-negative cells were systematically identified and excluded.

Perturb-seq data were clustered using the Seurat package, with cell state annotations assisted through integration with multiome-snRNA-seq data via reciprocal PCA (RPCA) (Figure S4F, G). The Seurat-RPCA was selected over Seurat-CCA here due to the largely missing early populations in the Perturb-seq data and the large numbers of cells. The NC (neural crest) subset of the Perturb-seq data was then isolated and subjected to further clustering. Cell type annotations were established based on the RPCA integration outcomes between the Perturb-seq NC and multiome-snRNA-seq NC datasets. Clusters with low feature counts were omitted from subsequent analyses. To minimize technical discrepancies, all knockout panels were sequenced to achieve relatively uniform depths (Figure S4E).

#### Quantification of knockout effects

The filtered Perturb-seq NC data were visualized using the PHATE method^189^. To measure the extent of gene disruption, the MELD method^72^ was used to quantify the differences in abundance between the control (Cherry no-targeting) and each knockout condition/panel. Given that each knockout condition encompassed multiple embryos (n ≥ 12), the Perturb-seq data reflected an average effect, thereby minimizing inter-animal variance. While the control and some conditions involved cells from several 10x sequencing runs, aiding in mitigating batch effects, technical duplicates were still produced through re-sampling to compute a mean MELD likelihood value.

To gauge the short-term impacts of the knockout, the latent time for the affected cells was determined. Two integration approaches, RPCA^190^ and scVI^191^, were employed to generate shared-nearest-neighbor graphs. The average latent time of neighboring cells was computed for each cell, serving as the imputed latent time. The latent times estimated via RPCA and scVI demonstrated a significant correlation (Figure S4H), and the RPCA-estimated latent time was selected for further downstream analysis and visual representation. For each lineage, the median of estimated latent time was used to dichotomize cells in this lineage-specific cluster into early and late stages. Finally, the depletion-enrichment likelihood (range: 0-1) was computed by MELD, and the average likelihood of the late-stage cells for each lineage represented the lineage-specific short-term effect for each knockout panel.

#### Differential expression analysis

To identify differentially expressed genes between control and knockout cells, we performed cluster-specific differential expression analysis, where cells within each cluster were randomly aggregated into pseudo-bulk duplicates by the R function Matrix.utils::aggregate.Matrix. For each cluster, edgeR analysis^192^ was then conducted with the functions calcNormFactors, estimateDisp, glmQLFit, and glmQLFTest to compare control and a certain knockout condition.

## Data availability

All raw data, processed data, and metadata from the 10x Multiome, MERSCOPE, direct capture perturb-seq, ChIP-seq, and genotyping amplicon-seq have been deposited to the Gene Expression Omnibus (accession: GSE256011). The Smart-seq3 dataset from ref^43^ can also be obtained from GSE256011. The 10x Multiome and MERSCOPE data are available to view on the UCSC Cell browser (https://zhiyhu.github.io/neural_crest_cellbrowser/). The interactive zebrafish NC-GRN viewer with options to select regulons and view their downstream targets is available at https://zhiyhu.github.io/NC-GRN_shiny/.

## Code availability

All analysis code is available at the GitHub repository (https://github.com/zhiyhu/neural-crest-scmultiomics).

---

**Supplementary Figure 1.**
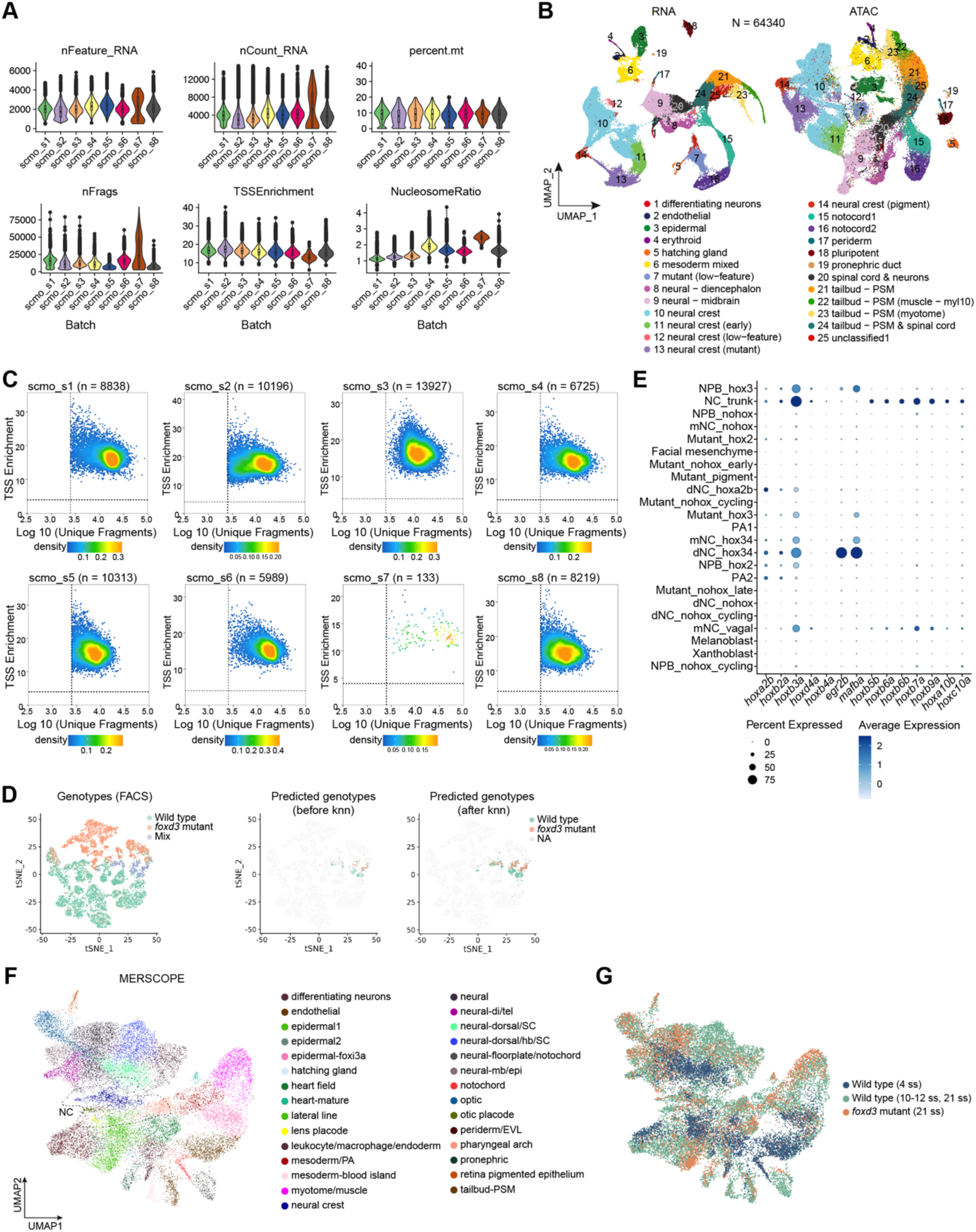
Quality control and cell type overview across 10x Multiome and spatial transcriptomics datasets. Related to Figure 1. **(A)** Violin plots show uniform quality across samples in the 10x multiome data indicated by metrics, namely the number of detected genes per cell (nFeature_RNA), RNA UMI counts per cell (nCount_RNA), mitochondrial gene percentage (percent.mt), ATAC fragment count (nFrags), transcription start site enrichment (TSSEnrichment), and nucleosome ratio (NucleosomeRatio). The upper and lower edges of the boxplots indicate the 25% and 75% quantiles, and the middle line shows the median. **(B)** UMAP plots show that cell type annotations are reliably reflected by both multiome-snRNA-seq and multiome-snATAC-seq datasets. **(C)** The quality of filtered 10x multiome-snATAC-seq data is demonstrated by the log10 of unique fragments and TSS enrichment, with optimal samples densely distributed in the second quadrant. Dashed lines represent quality thresholds. **(D)** *t-*SNE plots visualizing the results of computational demultiplexing, with cells colored according to ground-truth genotypes, logistic regression model predictions, and *k-*nearest neighbor (*k*-NN) refinement. **(E)** HOX family gene and position-specific marker expression patterns, facilitating the cranial-caudal axial-level cell state annotation. **(F)** Cell type annotations of the MERSCOPE dataset. PA, pharyngeal arch; SC, spinal cord; hb, hindbrain; mb, midbrain; epi, Epiphysis; EVL, enveloping layer; PSM, pre-somite mesoderm. **(G)** Sample annotations in the MERSCOPE dataset of wild-type and *foxd3*-mutant embryos.

**Supplementary Figure 2.**
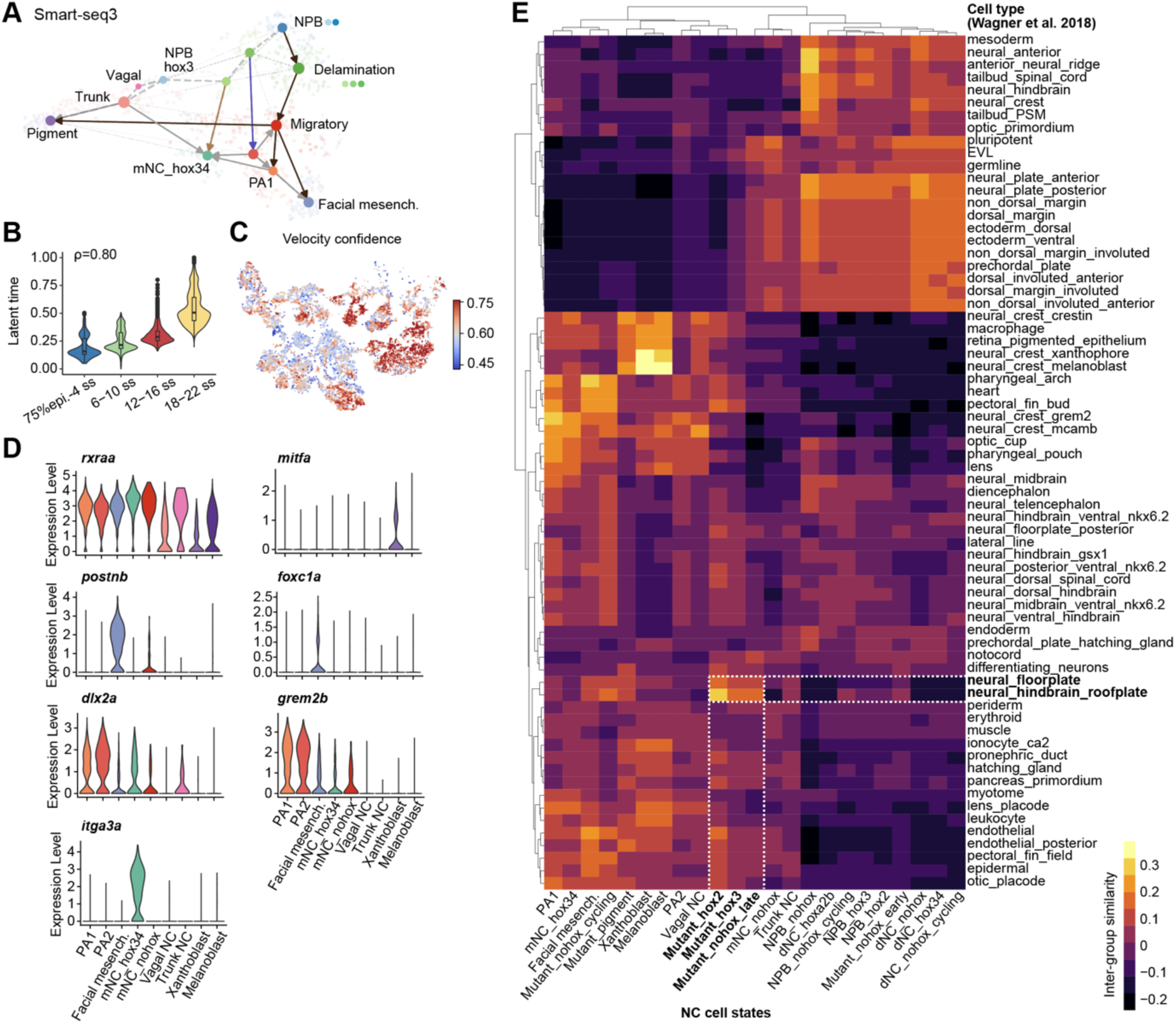
Validation of cranial NC trajectories and cell states. Related to Figure 2. **(A)** Developmental trajectories of wild-type NC reconstructed using full-length transcript Smart-seq3 data. **(B)** Boxplot shows a positive correlation of latent time with developmental stages, labelled by the Spearman correlation coefficient. **(C)** *t*-SNE plot shows the velocity confidence of wild-type NC cells. **(D)** Violin plots show the expression patterns of marker genes validated in Figure 2C. **(E)** The inter-group similarity heatmap computed by CIDER^173^ shows the resemblance of *foxd3*-mutant NC cells and floorplate or roofplate cells in terms of differential gene expression patterns.

**Supplementary Figure 3.**
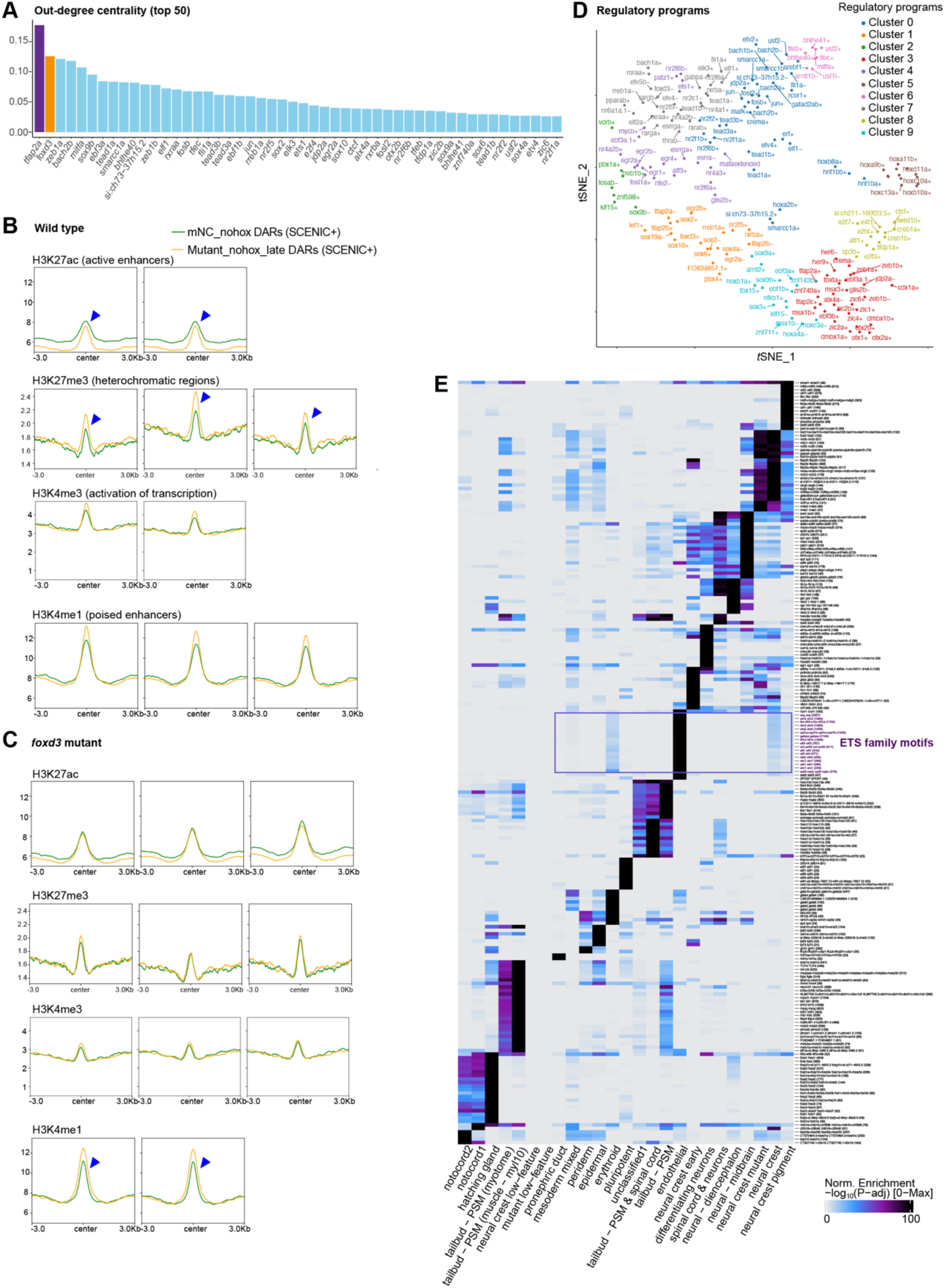
Functional analysis of the SCENIC+-predicted NC GRN. Related to Figure 3. **(A)** Out-degree centrality analysis identifies *tfap2a* and *foxd3* as pivotal regulators in the SCENIC+-inferred NC GRN. **(B-C)** MicroChIP-seq profiles of enhancer candidates (i.e., differentially accessible regions, DARs) for two clusters (mNC_nohox and Mutant_nohox_late) in two samples, namely *foxd3*-expressing wild-type migratory cells **(B)** and in *foxd3*-mutant cells **(C)**. **(D)** *t*-SNE plot exhibits the results from agglomerative clustering of regulon region-based activities, with regulons colored by the regulatory programs (clusters). |**(E)** Motif enrichment analysis of multiome-snATAC-seq data highlights that ETS motifs are enriched in wild-type NC cells and absent in *foxd3* mutants.

**Supplementary Figure 4.**
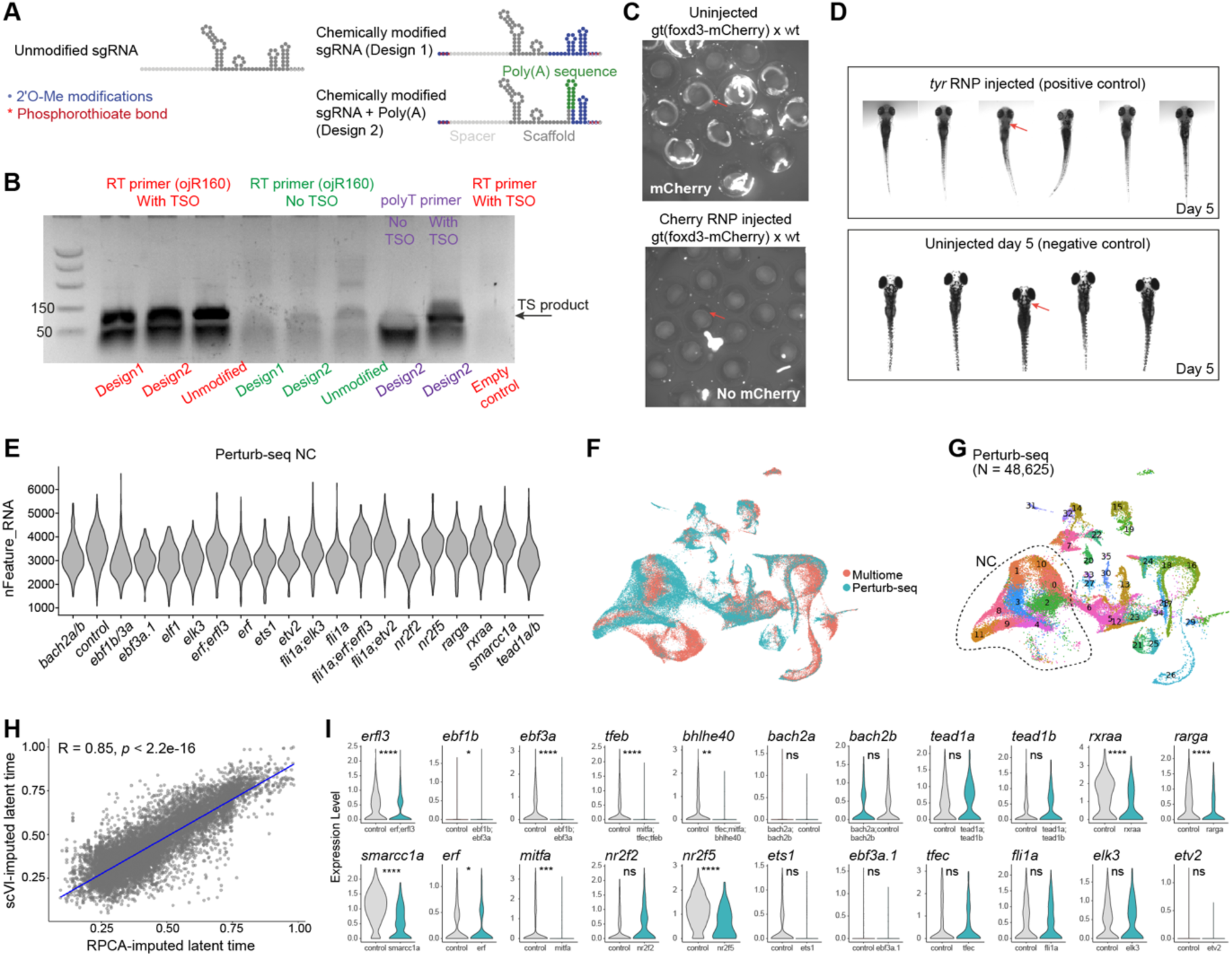
sgRNA design and knockout efficacy of chemically modified sgRNAs and Perturb-seq data overview. Related to Figure 4. **(A)** Schematic representations of unmodified, chemically modified, and chemically modified plus poly(A) sgRNA structures. **(B)** Gel electrophoresis confirms that chemical modifications do not impair the template-switching reaction in the scRNA-seq protocol. TS, template switching reaction; RT, reverse transcription; TSO, template switching oligo. The DNA ladder size (50bp and 150bp) is labelled. **(C)** Contrast images validate the elimination of mCherry fluorescence (indicated by red arrows) in gt(foxd3-mCherry) x wild-type embryos following RNP injection with three sgRNAs (Design 1) targeting the mCherry sequence. **(D)** Contrast images exhibit reproducible phenotypes in *tyr* F_0_ crispants on day 5 post injection, with red arrows highlighting the phenotypes. **(E)** Violin plot displays uniform gene coverage across knockout conditions. **(F)** UMAP visualization presents the integrated landscape of multiome-snRNA-seq and Perturb-seq data. **(G)** Perturb-seq data clusters are indicated within the same integrated space as (F). NC clusters were identified based on multiome cell annotations. **(H)** Scatterplot confirms the significant correlation between latent time predictions using RPCA and scVI. **(I)** Violin plots show gene expression levels for targeted TFs in non-targeting control versus knockout conditions. N.s., not significant; *, *p* < 0.05; **, *p* < 0.01; ***, *p* < 0.001; ****, *p* < 0.0001, by Wilcoxon rank sum test.

**Supplementary Figure 5.**
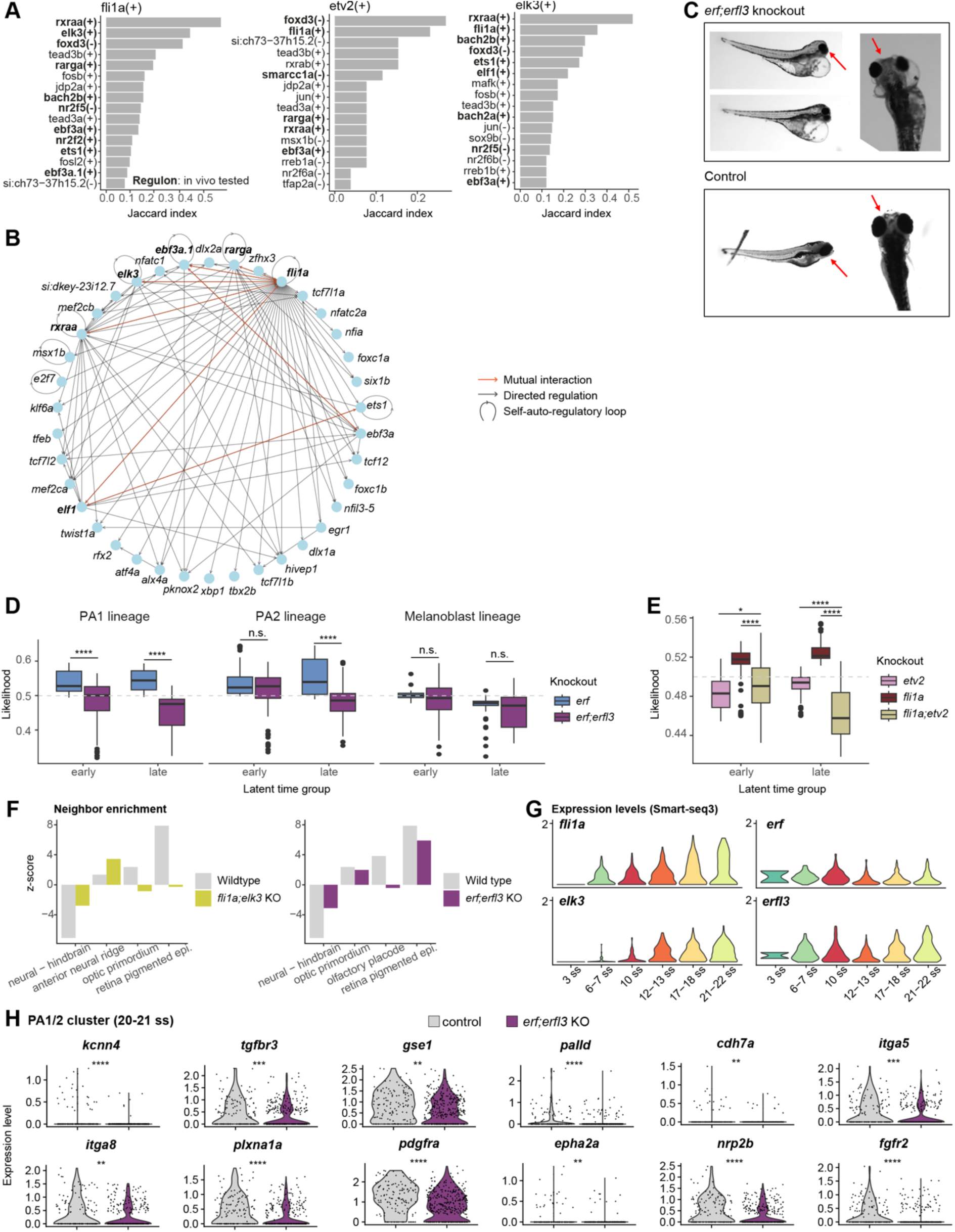
Individual and convergent functions of ETS TFs in NC development. Related to Figure 5. **(A)** Bar plots display notable functional redundancy among ETS TFs (*etv2, fli1a, elk3*), as evidenced by their regulons sharing the highest Jaccard indices. A high degree of target-gene overlap with *foxd3* and RXR/RAR/NR2F regulons is observed for several ETS TFs (*elk3, fli1a, ets1*), suggesting potential synergistic effects. **(B)** Ego graph from the SCENIC+-predicted GRN for *fli1a* demonstrates self-auto-regulatory loops and mutual regulatory interactions between *fli1a* and *elk3*, as well as between *fli1a* and *rxraa/rarga* indicating intricate regulatory networks of ETS TFs. **(C)** Phenotypic analysis of *erf;erfl3* double knockout embryos indicates significant craniofacial abnormalities (highlighted by red arrows). **(D)** In vivo Perturb-seq analysis shows profound depletion effects for *erf;erfl3* double-TF knockouts compared to *erf* single-TF knockouts in late PA1 and PA2 lineages. ****, *p* < 0.0001; n.s., not significant, by Wilcoxon rank sum test. **(E)** In vivo Perturb-seq analysis shows profound depletion effects for *fli1a;etv2* double-TF knockouts compared to *fli1a* or *etv2* single-TF knockouts in the late non-hox facial mesenchymal lineage. ****, *p* < 0.0001; *, *p* < 0.05, by Wilcoxon rank sum test. **(F)** Spatial neighbor enrichment analysis results comparing the wild-type and knockout embryos. The *z*-score is computed using the permutations test based on the cell cluster proximity. A higher *z*-score represents a cell type enrichment in the neighbor cells. The neural-hindbrain and the anterior neural ridge clusters represent the dorsal cell populations, and the optic primordium, retina pigmented epithelium (epi.), and olfactory placode represent distal sites away from the NC delamination positions. **(G)** Expression patterns of representative ETS TFs across developmental stages within Smart-seq3 data. The y-axis indicates expression levels, and the x-axis indicates developmental stages. **(H)** Differentially expressed genes between control and *erf;erfl3-*knockout cells in the PA1/2 cluster. ****, *p* < 0.0001; ***, *p* < 0.001; **, *p* < 0.01; n.s., not significant, by edgeR.

**Supplementary Figure 6.**
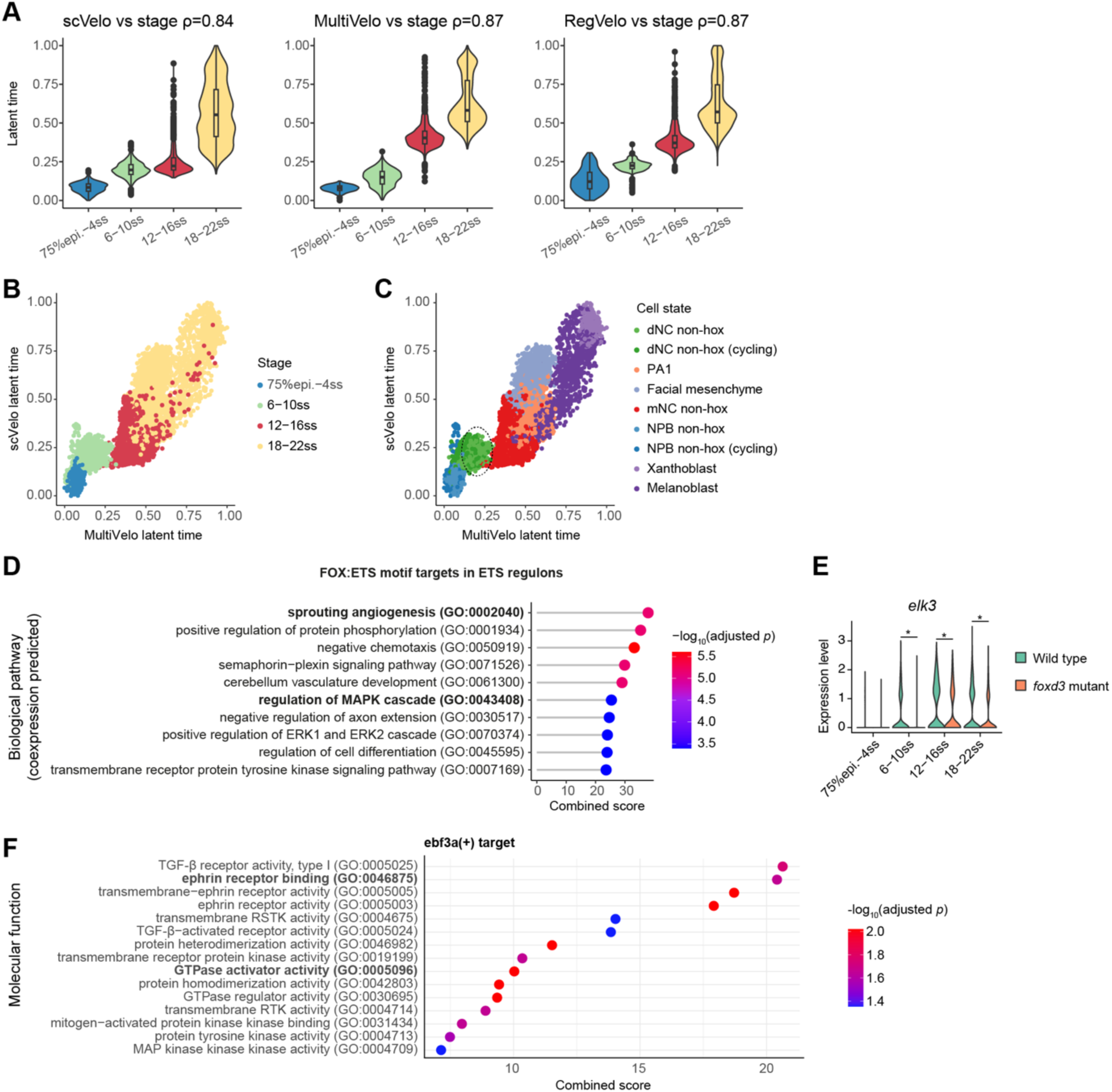
GRN temporal dynamics analysis of the endothelial-like migratory program. Related to Figure 6. **(A)** Relationship between chronological development stages and latent time predicted by scVelo, MultiVelo, and RegVelo, revealing that scVelo does not differentiate between dNC (6-10 ss) and mNC (12-16 ss) cell states, although all latent time prediction results show a positive association. The Spearman correlation coefficients (ρ) are indicated. Epi., epiboly. **(B-C)** Comparative evaluation of scVelo and MultiVelo latent times, with cells distinguished by developmental stages (B) and cell states (C). The dashed circle indicates the overlap between dNC and mNC in RNA-velocity-based latent time. **(D)** Gene ontology enrichment analysis for target genes of FOX:ETS motif in ETS regulons. The x-axis represents the FishEnrichR combined scores, and the y-axis indicates the gene ontology groups. **(E)** Downregulation of *elk3* in *foxd3*-mutant NC cells across developmental stages. The star indicates Benjamini-Hochberg (BH)-corrected *p* < 4×10^-28^, by limma voom^174^. **(F)** Gene ontology enrichment analysis for target genes of ebf3a(+) regulon.

**Supplementary Figure 7.**
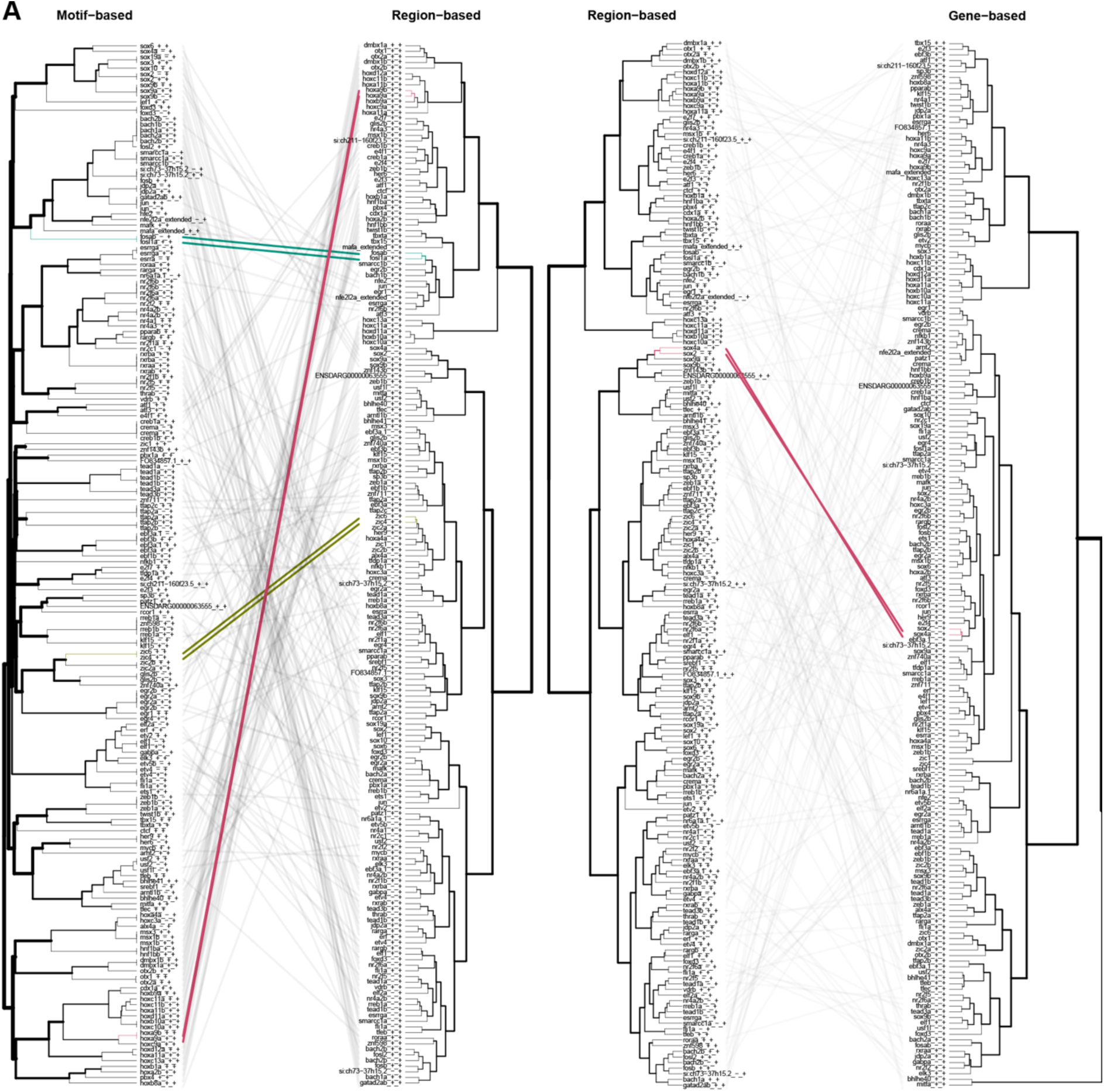
Regulon functional similarity comparison based on region activity, gene expression, and motifs. Related to Figure 7. **(A)** Comparative analysis of similarity metrics computed from motifs, region-based regulon dynamics, and gene-based regulon dynamics. The dendrograms of regulons were constructed based on one of the similarity metrics (referenced in subtitles), with gray lines illustrating connections between identical regulons across two dendrograms. Colored lines underscore the shared nearest neighbors between the two dendrograms.

**Supplementary Table 1.** Sample information of single-cell sequencing and spatial transcriptomics, related to Figure 1, Figure 4, STAR Methods.

**Supplementary Table 2.** Marker genes of NC cell states, related to Figure 1.

**Supplementary Table 3.** Cell-type-specific regulon specificity scores of NC GRN, related to Figure 3.

**Supplementary Table 4.** Biological processes identified by the gene ontology enrichment analysis conducted on the target genes of each functional cluster, related to Figure 3.

**Supplementary Table 5.** Gene ontology enrichment analysis of BACH, EBF and TEAD regulons, related to Figure 6.

**Supplementary Table 6**. Primer sequences and sgRNA spacer sequences, related to STAR Methods.

